# Inoculation with the mycorrhizal fungus *Rhizophagus irregularis* modulates the relationship between root growth and nutrient content in maize (*Zea mays* ssp. *mays* L.)

**DOI:** 10.1101/695411

**Authors:** M. Rosario Ramírez-Flores, Elohim Bello-Bello, Rubén Rellán-Álvarez, Ruairidh J. H. Sawers, Víctor Olalde-Portugal

## Abstract

Plant root systems play an essential role in nutrient and water acquisition. In resource-limited soils, modification of root system architecture is an important strategy to optimize plant performance. Most terrestrial plants also form symbiotic associations with arbuscular mycorrhizal fungi to maximize nutrient uptake. In addition to direct delivery of nutrients, arbuscular mycorrhizal fungi benefit the plant host by promoting root growth. Here, we aimed to quantify the impact of arbuscular mycorrhizal symbiosis on root growth and nutrient uptake in maize. Inoculated plants showed an increase in both biomass and the total content of twenty quantified elements. In addition, image analysis showed mycorrhizal plants to have denser, more branched root systems. For most of the quantified elements, the increase in content in mycorrhizal plants was proportional to root and overall plant growth. However, the increase in boron, calcium, magnesium, phosphorus, sulfur and strontium was greater than predicted by root system size alone, indicating fungal delivery to be supplementing root uptake.

## INTRODUCTION

Plant productivity is typically limited by the availability of nutrients, in both natural and agricultural ecosystems. Under nutrient-poor conditions, plants have the capacity to modulate the architecture and functionality of their root system, potentially increasing nutrient uptake (Lynch, 1995). Beyond a general reallocation of resources from leaf to root, there are nutrient-specific changes in the development of primary (PR) and lateral (LR) roots, building a root system architecture (RSA) that may better optimize resource acquisition (López-Bucio et al., 2003; Osmont et al., 2007; Gruber et al., 2013). Nutrients are heterogeneously distributed in the soil, and plants respond to local concentration, allocating greater root production to regions of higher availability (Campbell et al., 1991; Farley and Fitter, 1999; Grossman and Rice, 2012). In addition, nutrient distribution varies across soil horizons: poorly-mobile nutrients such as phosphorus (P), potassium (K), magnesium (Mg) or calcium (Ca) are typically enriched in topsoil, while more mobile nutrients, such as nitrogen (N), are typically more abundant deeper in the soil (Rubio et al., 2003; Ho et al., 2004; Lynch and Brown, 2008; Postma et al., 2014; Rangarajan et al., 2018). As a consequence, researchers have distinguished RSAs optimized for topsoil versus deeper foraging (Lynch, 2019).

Root development is regulated by the combined action of internal developmental pathways and external environmental stimuli (Malamy and Ryan, 2001), conditioning both plasticity and intra- and inter-specific variation. These pathways are based on the action of plant hormones, signal receptors, transcription factors and secondary messengers, including Ca^2+^, nitric oxide and reactive oxygen species (Fukaki and Tasaka, 2009; Garay-Arroyo et al., 2012; Jung and McCouch, 2013; Schlicht et al., 2013; Zhang et al., 2015; Shahzad and Amtmann, 2017). Auxin works with cytokinin to regulate LR initiation (Aloni et al., 2006), and with gibberellin to modulate cell proliferation and elongation (Fu and Harberd, 2003). Strigolactones impact LR formation and root hair development in a dose-dependent manner (Koltai, 2011), and, in maize, promote nodal root development (Guan et al., 2012). RSA is largely a product of the balance between root elongation and branching (Postma et al., 2014). Primary root growth will continue if the root meristem is active and populations of stem cells within the quiescent center are maintained. In *Arabidopsis*, the exhaustion of the primary root meristem under low P is considered the classic example of a plastic response to optimize topsoil foraging (Williamson et al., 2001; López-Bucio et al., 2002; Mora-Macías et al., 2017). Root branching, through the production of first- or higher-order LRs, is regulated at the level of the formation of LR primordia and their subsequent expansion. Low availability of N, P, sulfur (S) or zinc (Zn) promotes an increase in the density and/or length of LRs, although, as mentioned above, when the nutrient distribution is patchy, LRs may proliferate in regions of high local nutrient abundance (Zhang et al., 1999; Kutz et al., 2002; López-Bucio et al., 2002; Bouranis et al., 2008; Gruber et al., 2013). In cereal crops, the bulk of the adult root system is comprised of shoot-borne crown (CR) roots and associated LRs (Hochholdinger and Tuberosa, 2009). While greater branching will increase the surface area for nutrient uptake, it becomes inefficient if placement of roots in close proximity results in competition for the same nutrients, an effect that will be greater for more mobile nutrients (Postma et al., 2014).

Significant intra-specific variation has been observed in RSA and plasticity, and has been linked to superior agronomic performance in crops under specific edaphic conditions. In common bean, varieties producing shallow basal roots and a large number of adventitious roots explore the topsoil more efficiently, performing better in low P soils (Rubio et al., 2003; Lynch and Brown, 2008). In rice, the PHOSPHATE STARVATION TOLERANCE 1 (PSTOl1) kinase, identified from the traditional variety Kasalath, is associated with enhanced early root growth, and increases yield under low P conditions (Wissuwa et al., 2005; Gamuyao et al., 2012).

In addition to relying on their roots to acquire nutrients, most terrestrial plants also form mutualistic symbioses with arbuscular mycorrhizal (AM) fungi of the Phylum Glomeromycota (Parniske, 2008; Smith and Read, 2010). The association with AM fungi is one of the oldest plant symbioses, and AM symbioses were likely important during plant colonization of the land, performing essential functions before the evolution of the vascular root system (Schüßler et al., 2001). Establishment of AM symbiosis requires a complex interchange of signals between plant and fungus that results in the entry of the fungus into the root and the development of highly branched arbuscules within the root cortical cells, providing the primary site of nutrient exchange (Parniske, 2008). AM fungi are obligate symbionts, obtaining carbon from their plant host in return for providing nutrients and water that are acquired by an extensive network of root-external hyphae (Bago et al., 2003). The clearest benefit to the plant is enhanced P uptake (Chiu and Paszkowski, 2019), although AM fungi have been reported to promote uptake of other elements, including N, Fe, S and Zn (Liu et al., 2000; González-Guerrero et al., 2005; Govindarajulu et al., 2005; López-Pedrosa and González-Guerrero, 2006; Allen and Shachar-Hill, 2009). In practice, complex interactions between nutrients influence the outcome of the symbiosis with respect to any given element (Liu et al., 2000; Gerlach et al., 2015; Ramírez-Flores et al., 2017). Although AM symbioses are widespread in both natural ecosystems and cultivated fields, the plant host maintains a degree of control over establishment and the extent of colonization, rejecting the fungus under high nutrient conditions (Nouri et al., 2014).

AM fungi not only directly deliver nutrients, but also promote growth of the host plant roots, with secondary effects on nutrient uptake. General improvements in plant health under AM symbiosis may be correlated with increased root system size (Berta et al., 1995; Tisserant et al., 1996; Gutjahr et al., 2009; Sawers et al., 2017). AM symbiosis is also associated with developmental changes, such as an increase in LR production and greater root branching (Berta et al., 1990; Paszkowski and Boller, 2002; Oláh et al., 2005; Gutjahr et al., 2009). The failure to form LR primordia in the maize *lateral rootless1* (*lrt1*) mutant can be partially overcome by AM symbiosis, indicating an influence on plant developmental pathways (Hochholdinger and Feix, 1998; Paszkowski and Boller, 2002). In *Medicago truncatula*, developing fungal spores are sufficient to trigger LR formation (Oláh et al., 2005; Gutjahr et al., 2009). In rice, AM fungi can induce LR formation in *pollux*, *ccamk* and *cyclops* mutants, even though symbiosis is not established, confirming that RSA modification need not be dependent on enhanced host plant nutrition (Gutjahr et al., 2009).

The impact of AM symbiosis on nutrient uptake can be quantified by comparing the concentration or content of a given element in the aerial portion of mycorrhizal (M) and non-colonized (NC) plants (Gerlach et al., 2015; Ramírez-Flores et al., 2017). Such a comparison, however, does not distinguish direct hyphal nutrient delivery from the secondary consequences of fungus-induced modification of root architecture or the effects of greater root growth resulting from a general increase in plant vigor. In the case of P, elegant labelling experiments and extensive functional studies have clearly demonstrated the direct role of root-external hyphae in nutrient delivery (Pearson and Jakobsen, 1993; Smith et al., 2003; Chiu and Paszkowski, 2019). The picture, however, remains less clear with respect to other mineral nutrients.

In this study, we characterized RSA and leaf ionome in young maize plants, grown with or without inoculation with AM fungi. For each nutrient, we asked whether any increase in uptake was explainable by changes in the root system alone, or whether there was evidence of additional hyphal foraging. For an increase resulting from greater root growth, we expected the relationship between root surface area and uptake to be maintained, albeit that mycorrhizal plants themselves were larger. If, however, there was a significant effect of hyphal foraging, we predicted an increase in uptake per unit surface area of root.

## MATERIALS AND METHODS

### Plant material and growth conditions

Two maize inbred lines (*Zea mays* ssp. *mays* var. B73 and W22) were grown with (M) or without (NC) inoculation with fungus, in PVC tubes (1 m in height, 15 cm in diameter). A total of 64 plants (2 genotypes x 2 treatments x 16 replicates) were planted in complete blocks across four planting dates. Each planting consisted of 16 plants (2 genotypes x 2 treatments x 4 replicates), with a one week interval between planting dates. B73 seed was produced in the winter of 2015 in Valle de Banderas, Nayarit, México. W22 seed was produced in the summer of 2013 in Aurora, New York, USA. Each tube was filled with 17 L of a sterilized substrate mix consisting of sand:perlite:silt (4.5:1.5:1, v/v). The substrate Olsen P concentration was 4.8 ppm. For M plants, we inoculated each tube with ~700 *Rhizophagus irregularis* spores collected from a commercial liquid inoculant (AGTIV^®^) and delivered to the middle of the tube at 15 cm depth before planting. The experiment was conducted under greenhouse conditions, at an average temperature of 24 °C and humidity of 48%. Maize seeds were surface sterilized with 1% sodium hypochloride solution for 10 min and rinsed five times with sterile distilled water. Seeds were then placed in 30 ml of sterile distilled water and shaken for 48 hours before planting. From 5 days after emergence (DAE), plants were watered every other day with 200mL of ⅓ Hoagland solution, modified to a final P concentration of 25 μM (Hoagland and Broyer, 1936). Plants were harvested at 56 DAE.

### Evaluation of plant growth

Leaf length and width were measured manually, and leaf area estimated as 0.75 x length x width (Mollier and Pellerin, 1999). Shoot and root fresh weights were measured after cutting the aerial part and washing the root. Dry weight was determined by oven drying tissue at 70° for 48 hrs. Root volume was measured by submergence in a test tube and recording of the volume of water displaced.

### Estimation of fungal colonization in seedling roots

Lateral root segments were collected at random from the upper 15 cm of the root system, placed in 10% KOH solution, autoclaved for 10 min, and stained with 0.05% trypan blue in acetoglycerol. The percentage of root length colonized was quantified in 15 root pieces per plant using a modified grid-line intersect method (McGonigle et al., 1990).

### Characterization of root system architecture by image analysis

The cleaned root system was placed in a water-filled tub and photographed using a digital Nikon camera D3500. Raw images were individually processed using Adobe^®^ Photoshop^®^ CC (Version 14.0) to remove background and increase contrast. Processed images were scaled and analyzed using GiA Roots software (Galkovskyi et al. 2012). After scale calibration, the images were segmented using the adaptive thresholding method and basic parameter settings were manually optimized. Finally, all binary images were analyzed to quantify the root system traits. Stacked (overlay) root system images were generated using the stacks tool in ImageJ/Fiji (version 2.0.0). Root traits are described in Supplementary Table S1.

### Analysis of the leaf ionome

The third youngest leaf (variously, leaf 5 or 6 for different individuals) was collected for ionomic analysis using inductively coupled plasma mass spectrometry (ICP-MS) as described previously (Ziegler et al. 2013, Ramírez-Flores et al., 2017). Briefly, weighed tissue samples were digested in 2.5 ml of concentrated nitric acid (AR Select Grade, VWR) with an added internal standard (20 p.p.b. In, BDH Aristar Plus). The concentration of the elements B, Na, Mg, Al, P, S, K, Ca, Mn, Fe, Co, Ni, Cu, Zn, As, Se, Rb, Sr, Mo and Cd was measured using an Elan 6000 DRC-e mass spectrometer (Perkin-Elmer SCIEX) connected to a PFA microflow nebulizer (Elemental Scientific) and Apex HF desolvator (Elemental Scientific). A control solution was run every 10th sample to correct for machine drift both during a single run and between runs. Given that samples were digested in concentrated nitric acid prior to analysis and run in a 70% N atmosphere, it was not possible to accurately estimate N concentration.

### Statistical analyses and data visualization

All statistical analyses were performed in R (R Core Team, 2019). From the initial planting, a small number of plants did not germinate. In addition, we removed clear outliers (assessed by visual inspection of plant growth and health) and any plants assigned to the non-inoculated group that showed evidence of root-internal hyphal structures. The final dataset consisted of 53 individuals (Table S2). Leaf surface area estimates were square root transformed for analysis. A linear fixed-effect model was applied on a trait-by-trait basis to control for differences between the four planting dates. Principal component (PC) analysis was performed on Gia Roots traits along with root fresh (RFW) and dry weight (RDW), using R/ade4::dudi.pca (Dray and Dufour, 2007) with centered and scaled values. Linear Discriminant (LD) analysis on PC scores was performed with R/MASS::lda (Venables and Ripley, 2002). Ionomic analysis generated element concentration data; in addition, total element content was estimated as the product of concentration and shoot dry weight (SDW).

Data were analyzed using R/stats::lm to fit inoculation status and genotype in a complete model. After adjustment for multiple testing (R/stats::p.adjust; method = “BH”), no significant (p < 0.05) interaction between inoculation x genotype was detected for any trait, and we present here the additive model for simplicity. We also performed a one-way ANOVA with a single four-level treatment factor (B73.NC, B73.M, W22.NC and W22.M) that was used to assign means groups with R/agricolae::HSD.test (de Mendiburu, 2019) for presentation in Fig. S1. Box plots were generated with base R using default settings. Root images were pre-processed using imagemagick (www.imagemagick.org), converting them to png format, setting the background to transparent and reducing the alpha level. Images were imported into R using R/png and plotted onto the PCA/LD space using the rasterImage function. Pearson’s correlation coefficients were calculated using R/Hmisc::rcorr (Harrell, 2019) and visualized using R/gplots::heatmap.2 (Warnes et al., 2019). The impact of root system size and inoculation on element content, was assessed using the model *Content* = *NSA* + *Inoculation*, evaluating the *Inoculation* term using sequential (Type I) sum-of-squares (implemented by default in R). For comparison, partial (Type III) sum-of-squares were calculated using R/car::Anova (Fox & Weisberg, 2011), setting the “contrasts” option to c(“contr.sum”, “contr.poly”).

## RESULTS

### Growth of maize seedlings increased when inoculated with *Rhizophagus irregularis*

To evaluate the relationship between root system architecture (RSA) and nutrient acquisition in mycorrhiza plants, we grew the two maize inbred lines B73 and W22 under low phosphorus (P) conditions, with (M) or without (NC) inoculation with the AM fungus *Rhizophagus irregularis* (Fig. 1A). These two widely used inbred lines were selected to provide a reference for future work, and to allow greater generalization than would be possible based on a single genotype. B73 is the reference for the most complete maize genome assembly. W22 is the background for the primary publicly available maize reverse genetics collections. Inoculation with AM fungus significantly enhanced plant growth in both genotypes (Fig. 1B, S1, S2. Table 1). Fungal inoculation and plant genotype were both significant (p < 0.001) predictors of plant biomass but there was no evidence of an interaction between the two (Fig. S2, Table 1; S2). By harvest (55 DAE), the marginal effect of inoculation on shoot dry weight (SDW) was an increase of 142% (Table 1). In both M and NC treatments, B73 tended to be smaller than W22, the range of SDW overlapping between B73 M and W22 NC treatments (Fig. 1, S1, S2). The inclusion of plants similar in size, yet differing in inoculation status, was important for our subsequent interpretation of element content.

**Table 1.**
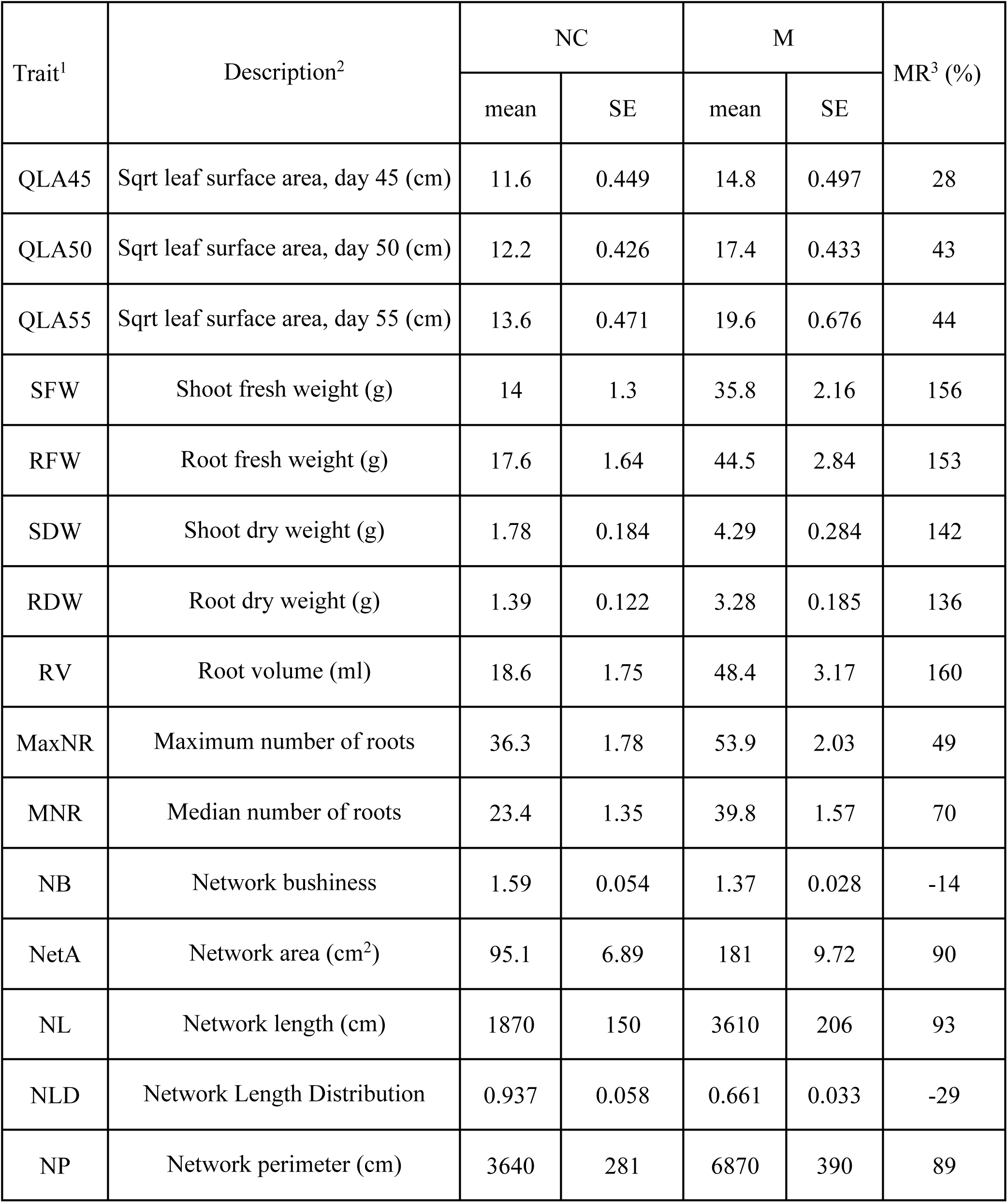
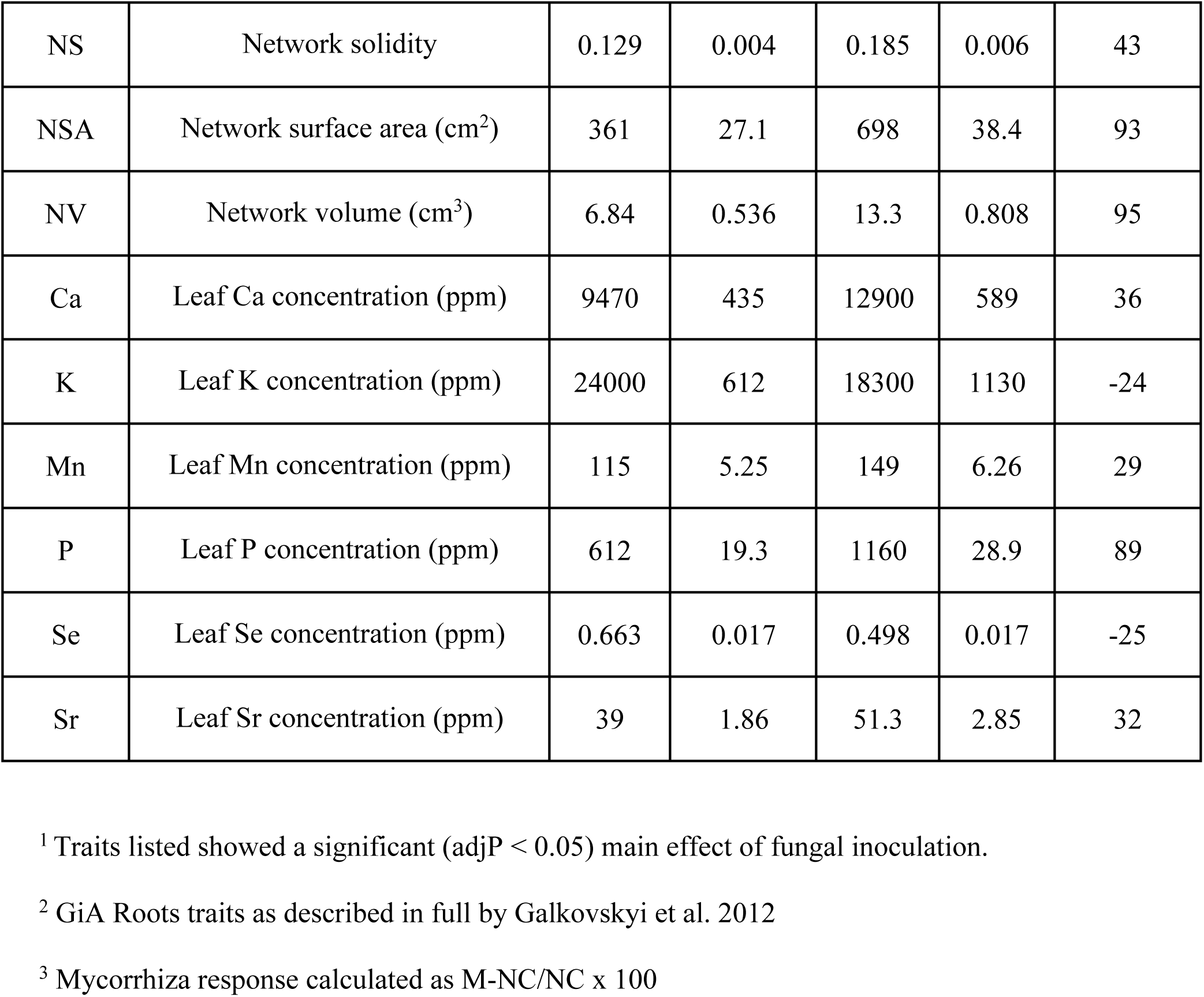
Traits responsive to inoculation with *Rhizophagus irregularis*

| Trait<sup>1</sup> | Description<sup>2</sup> | NC |  | M |  | MR<sup>3</sup> (%) |
| --- | --- | --- | --- | --- | --- | --- |
|  |  | mean | SE | mean | SE |  |
| QLA45 | Sqrt leaf surface area, day 45 (cm) | 11.6 | 0.449 | 14.8 | 0.497 | 28 |
| QLA50 | Sqrt leaf surface area, day 50 (cm) | 12.2 | 0.426 | 17.4 | 0.433 | 43 |
| QLA55 | Sqrt leaf surface area, day 55 (cm) | 13.6 | 0.471 | 19.6 | 0.676 | 44 |
| SFW | Shoot fresh weight (g) | 14 | 1.3 | 35.8 | 2.16 | 156 |
| RFW | Root fresh weight (g) | 17.6 | 1.64 | 44.5 | 2.84 | 153 |
| SDW | Shoot dry weight (g) | 1.78 | 0.184 | 4.29 | 0.284 | 142 |
| RDW | Root dry weight (g) | 1.39 | 0.122 | 3.28 | 0.185 | 136 |
| RV | Root volume (ml) | 18.6 | 1.75 | 48.4 | 3.17 | 160 |
| MaxNR | Maximum number of roots | 36.3 | 1.78 | 53.9 | 2.03 | 49 |
| MNR | Median number of roots | 23.4 | 1.35 | 39.8 | 1.57 | 70 |
| NB | Network bushiness | 1.59 | 0.054 | 1.37 | 0.028 | -14 |
| NetA | Network area (cm<sup>2</sup>) | 95.1 | 6.89 | 181 | 9.72 | 90 |
| NL | Network length (cm) | 1870 | 150 | 3610 | 206 | 93 |
| NLD | Network Length Distribution | 0.937 | 0.058 | 0.661 | 0.033 | -29 |
| NP | Network perimeter (cm) | 3640 | 281 | 6870 | 390 | 89 |
| NS | Network solidity | 0.129 | 0.004 | 0.185 | 0.006 | 43 |
| NSA | Network surface area (cm<sup>2</sup>) | 361 | 27.1 | 698 | 38.4 | 93 |
| NV | Network volume (cm<sup>3</sup>) | 6.84 | 0.536 | 13.3 | 0.808 | 95 |
| Ca | Leaf Ca concentration (ppm) | 9470 | 435 | 12900 | 589 | 36 |
| K | Leaf K concentration (ppm) | 24000 | 612 | 18300 | 1130 | -24 |
| Mn | Leaf Mn concentration (ppm) | 115 | 5.25 | 149 | 6.26 | 29 |
| P | Leaf P concentration (ppm) | 612 | 19.3 | 1160 | 28.9 | 89 |
| Se | Leaf Se concentration (ppm) | 0.663 | 0.017 | 0.498 | 0.017 | -25 |
| Sr | Leaf Sr concentration (ppm) | 39 | 1.86 | 51.3 | 2.85 | 32 |

<sup>1</sup> Traits listed showed a significant (adjP < 0.05) main effect of fungal inoculation.
<sup>2</sup> GiA Roots traits as described in full by Galkovskyi et al. 2012
<sup>3</sup> Mycorrhiza response calculated as M-NC/NC x 100

**Figure 1.**
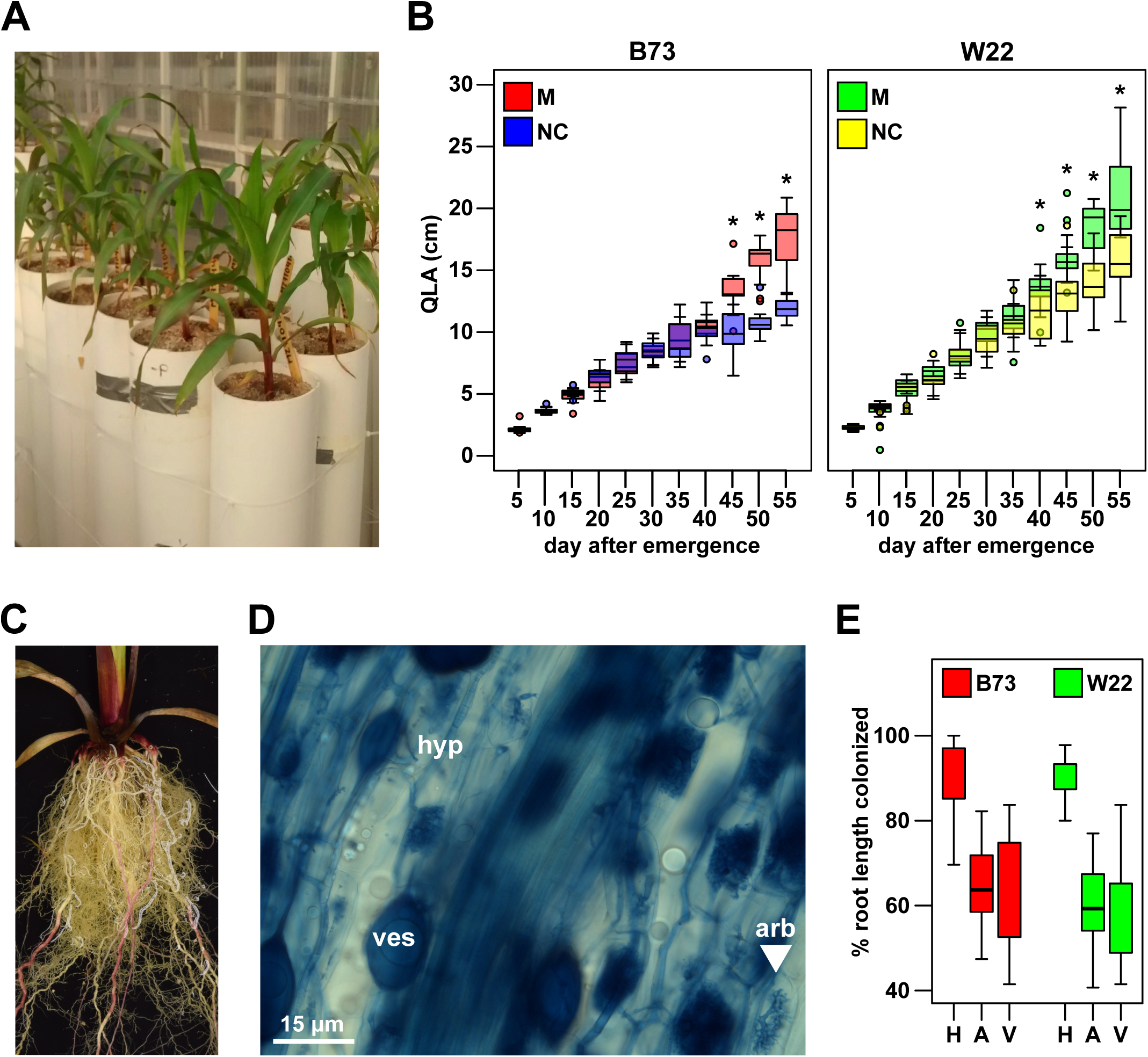
Experimental set-up and plant growth response. The maize inbred lines B73 and W22 were grown with (M) or without (NC) inoculation with the AM fungus *Rhizophagus irregularis*. A) Overall view of the growth system, the block in the foreground are inoculated plants. B) Square root of leaf surface area (QLA) for M and NC individuals, quantified every 5 days from 5 days after emergence until harvest at day 55. Boxes show 1st quartile, median and 3rd quartile. Whiskers extend to the most extreme points within 1.5x box length; outlying values beyond this range are shown as filled circles. Days at which M and NC groups were significantly different for a given inbred (Tukey HSD; p < 0.05) are indicated with an asterisk. C) Root crown on an M plant, illustrating the profusion of lateral roots and characteristic yellow pigmentation. D) Trypan-blue stained root section of an M plant, showing root-internal hyphae (hyp), vesicles (ves), and arbuscules (arb). Scale bar, 15 μm. E) Colonization (% root length colonized) for B73 and W22, determined with respect to fungal hyphae (H), arbuscules (A) and vesicles (V). Points indicate individual plants. Boxes and whiskers as in B).

To quantify mycorrhizal colonization, the percentage of root length containing different fungal structures was estimated by microscopic inspection. NC plants were confirmed to be free from colonization (fungal structures were observed in three plants in the NC group; these individuals were not included in the analysis), while inoculated plants showed fungal structures typical of the symbiosis (hyphae, arbuscules and vesicles). M plants were well colonized (Fig. 1C-E), and there was no difference in colonization rate between the two plant genotypes (Fig S1. Table S2).

### The root system was modified by inoculation with *Rhizophagus irregularis*

Root fresh and dry weight (RFW and RDW) and root volume (RV) increased significantly in inoculated plants (Fig. 2A. Table 1). To characterize RSA, we photographed the plants and analyzed the images with GiA Roots (General Image Analysis of Roots; Galkovskyi et al., 2012). Ten GiA Roots traits showed a significant response to inoculation (Table 1. Fig. S1). In a principal component (PC) analysis using all 19 GiA Roots traits, the first four PCs explained 90% of the total variance (Fig. 2B; 2C). PC1 (explaining 59% of the total variance) was dominated by variables associated with overall root system and plant size (Fig. 2; 4A). PC2 (explaining 12% of the total variance) was associated with overall root system shape and aspect ratio (Fig. 2; 4A). PC3 (7%) and PC4 (7%) captured aspects of root branching and root system solidity (Fig. 3; 4A). NC and M plants were well differentiated by PC1 and, to a lesser extent, by PC3, indicating a shift towards larger, more branched/solid root systems in M plants. GiA Roots evaluated solidity as the total network area divided by the network convex area. Although, greater solidity was clear in M W22, the root system in the smallest B73 NC plants was also considered relatively solid because the convex area itself was so low. We performed a linear discriminant (LD) analysis using the root PC values, reinforcing the separation of M and NC plants by size and solidity of the root system and better distinguishing the W22 M and B73 NC treatments (Fig. 3).

**Figure 2.**
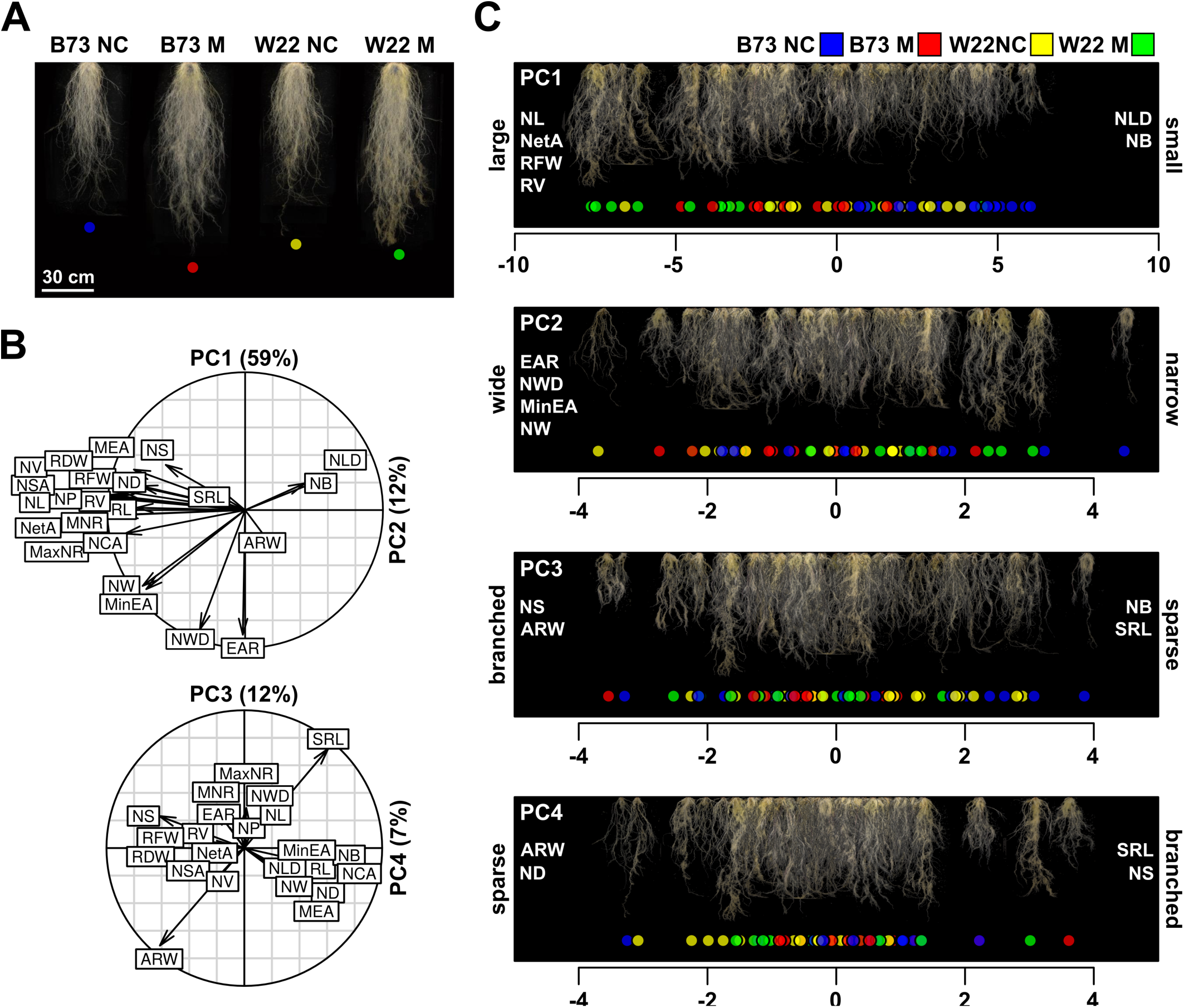
Principal component analysis describes variation in root system architecture. A) Stacked (overlay) images of the root systems of all plants in the NC/M B73/W22 treatment groups. Scale bar, 30 cm. B) The contribution of root system traits to principal components (PCs) 1 to 4. The variance explained by each PC is given in parentheses. See Table S1 for trait abbreviations. C) Root system images arranged by loading on each of PCs 1 to 4. Coloured points at the base of the image indicate genotype and fungal treatment as described in the key. Traits with a major positive or negative contribution to each PC are listed on the right or left of the plot, respectively. A single word description of the extremes of each PC is given at the side of each plot.

**Figure 3.**
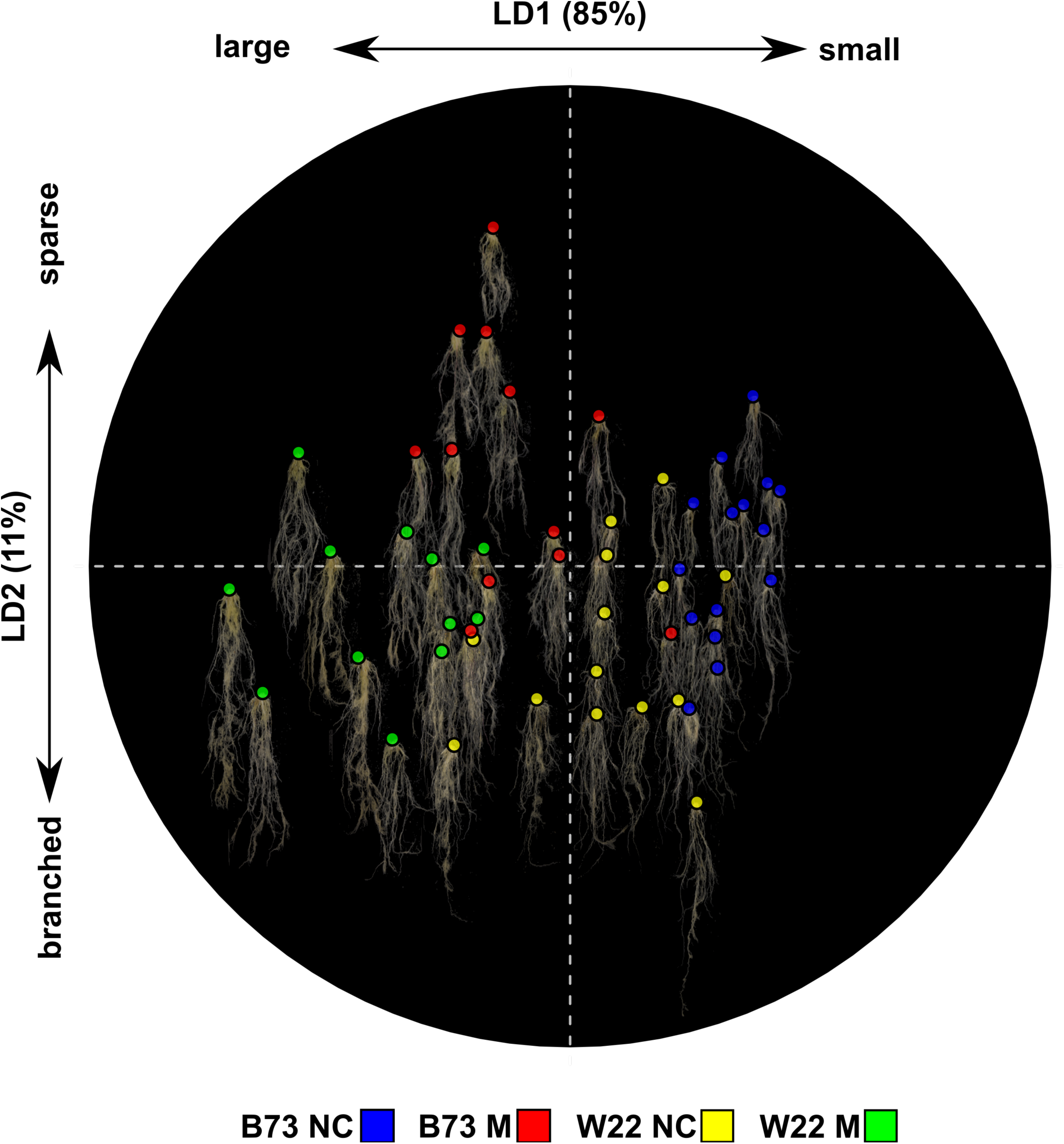
Inoculation with AM fungi is correlated with an increase in size and branching of the root system. Root system images projected by Linear Discriminants (LDs) 1 and 2. Colored points indicate genotype and fungal treatment as described in the key.

### Nutrient content was correlated with root system size and mycorrhizal colonization

AM fungi can increase plant nutrient uptake by delivery of nutrients via the hyphal network and as a secondary consequence of promoting root growth. To better understand the relative importance of these two factors, we considered the relationship between root system size and mineral nutrient content. We determined the concentration of 20 different elements, including P, in leaf tissue using inductively coupled plasma mass spectrometry (ICP-MS). Although leaf element concentration provides an endpoint readout of various aspects of plant nutrient relations (Salt et al., 2008), it was used here as a proxy for nutrient uptake. We estimated nutrient content as the product of leaf concentration and total shoot dry weight and investigated the relationship with the GiaRoots trait NSA (Network Surface Area), used as our measure of root system size. As expected, an increase in NSA (associated with larger plants) was correlated with greater nutrient content (Fig. 4B). Of greater interest, however, was the degree to which this relationship was modified by AM colonization; *i.e.* for any given nutrient, was NSA sufficient to explain element content in M plants? We ran a linear model using NSA and inoculation status as predictors, assessing inoculation based on sequential (Type I) sum-of-squares, evaluating any fungal effect beyond that explained by NSA alone (Fig. 5). A significant positive effect of inoculation was associated with boron (B), Ca, Mg, P, S and strontium (Sr), indicating a unit of root surface area to be associated with a greater leaf content of these elements in inoculated plants.

**Figure 4.**
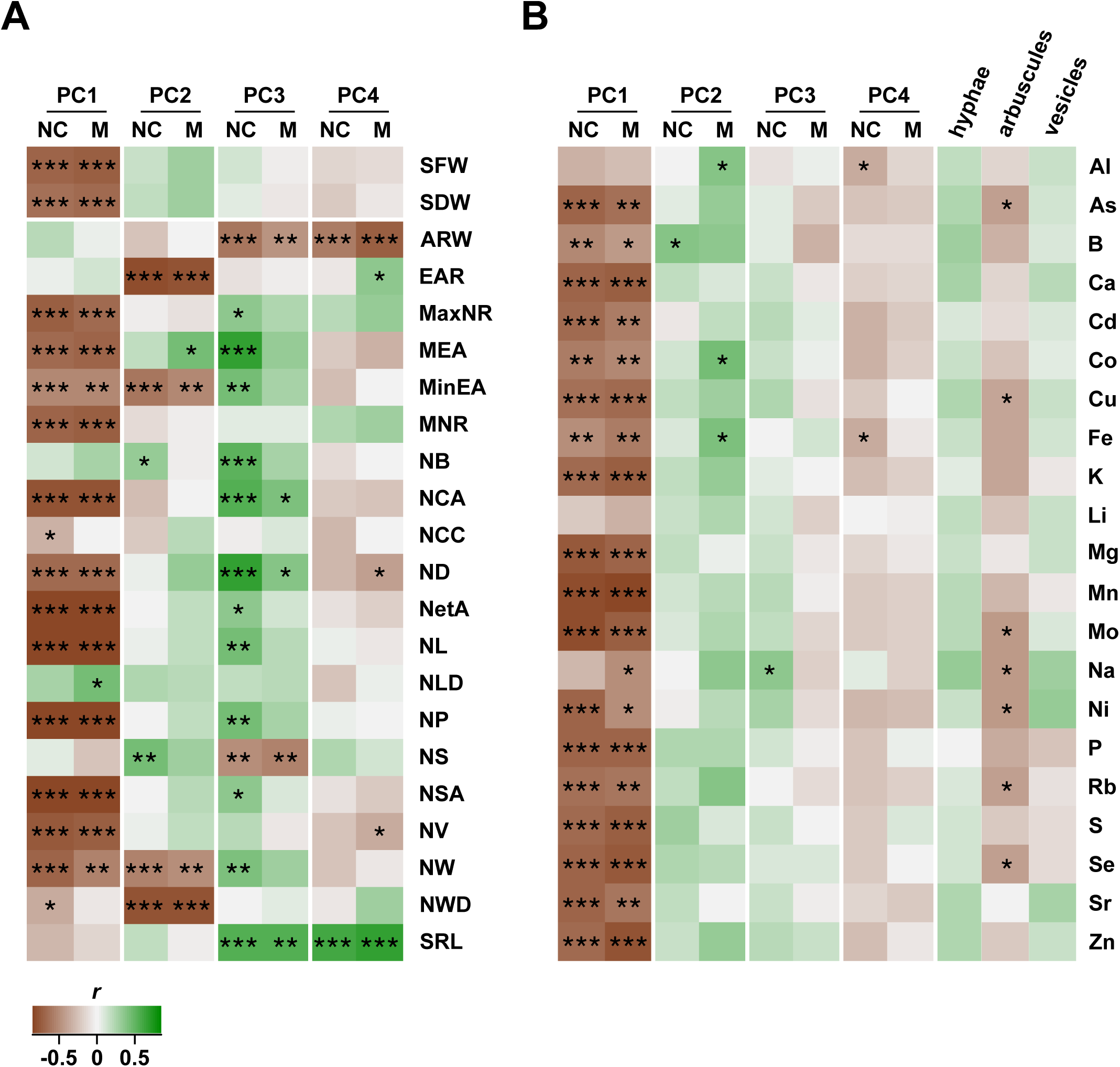
Root system architecture is correlated with total element content in the leaf. A) Heatmap representation of pairwise correlations of root system principal components (PC) with their contributing GiARoot traits (Abbreviated as described in Materials and Methods. SFW, shoot fresh weight and SDW, shoot dry weight also shown). B) Heatmap representation of pairwise correlations of root system principal components (PC) and colonization measures (% root length of hyphae (Hyp), vesicles (Ves) and arbuscules (Arb)) with total element content. In both A) and B) significant correlations are indicated by asterisks (*, p < 0.05; **, p < 0.01; ***, p < 0.001).

**Figure 5.**
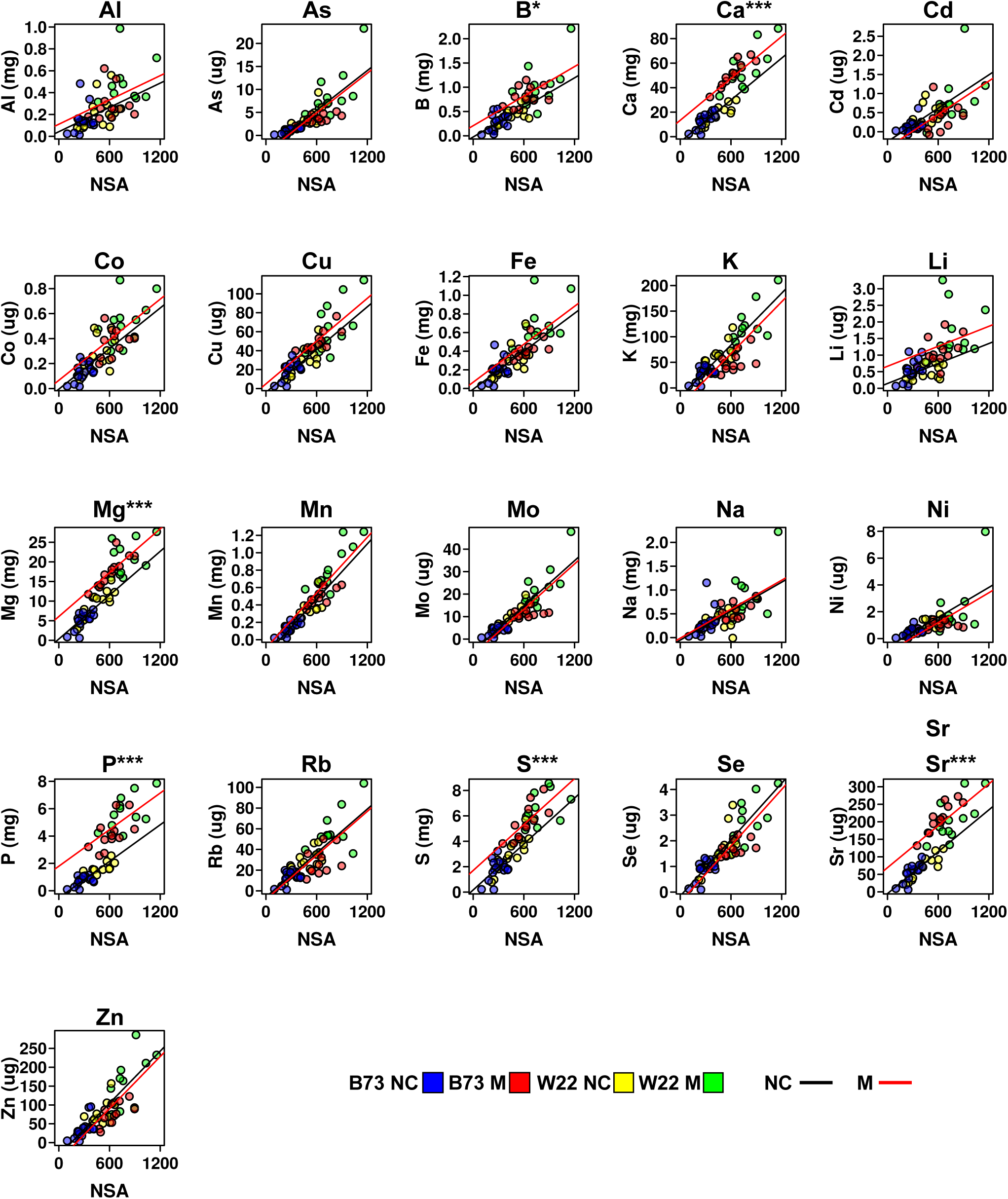
AM colonization modifies the relationship between root system size and element content. Scatter plot representation of the relationship between element content (shown in mg or μg) and root system size (estimated as the GiaRoots trait NSA) for twenty named elements. Colored points indicate genotype and fungal treatment of individual plants as described in the key. Black and red lines indicate the fit for the additive model *Total* ~ *NSA* + *Fungus* for levels NC and M of *Fungus*, respectively. The p-value for an effect of *Fungus* (calculated using Type I SS; adjusted for multiple tests) is given in parentheses following the element name in the plot title.

## DISCUSSION

Inoculation with *Rhizophagus irregularis* resulted in increased growth of maize grown under low P availability. In comparison with a previous small pot evaluation (Sawers et al., 2017), plants were larger, and the proportional increase in M plants was greater, presumably reflecting less growth inhibition in the larger tubes that we used here. A difference in leaf surface area was observed by day 40 - 45 after emergence, broadly consistent with the timing reported in rice for the first observation of arbuscules and the accumulation of transcripts encoding mycorrhiza-associated P transporters (Gutjahr et al., 2008). AM colonization was correlated not only with an increase in root system size, but also in the degree of branching and solidity of the root system (as captured by the GiA Roots traits MaxNR, MNR, NS). Inoculation with AM fungi has previously been shown to promote root growth and branching in diverse plant hosts (Berta et al., 1990; Paszkowski and Boller, 2002; Oláh et al., 2005; Gutjahr et al., 2009).

Increased growth was accompanied by greater concentration of Ca, Mn, P and Sr in the leaves of M plants. The leaf concentration of K and Se was reduced in M plants, but for all elements, taking biomass into account, total content increased. The content of B, Ca, Mg, P, S and Sr following inoculation exceeded expectation based on root surface area alone. We interpret this to reflect the impact of hyphal foraging, although we do not discount additional contributions of enhanced root function (*e.g.* greater density or length of root hairs; stimulation of the plant uptake pathways, production of exudates by plant or fungus) or changes in nutrient partitioning. For other elements, the content in M plants could be explained by the larger root system alone, although there is the possibility of a “hidden” mycorrhizal contribution (*i*.*e.* fungal nutrient delivery balanced by an equivalent reduction in direct root uptake. Smith et al., 2003).

The growth increase in inoculated plants was likely driven by increased P uptake, given that the experiment was conducted under low P availability. The route of P from soil to fungus, and subsequent delivery to the plant host is well characterized (Chiu and Paszkowski, 2019). We saw clear evidence that a unit of mycorrhizal root translates to a greater quantity of P obtained than an equivalent unit of non-colonized root, as would be anticipated as a consequence of hyphal foraging (Fig. 5). In comparison to P, the impact of AM colonization on the uptake of Ca, Mg and S (the other macronutrients for which our observations were consistent with hyphal foraging) is less well characterized (Gerlach et al., 2015; Ramírez-Flores et al., 2017). In the case of S, it has been shown that AM fungi have the capacity to transfer S from soil to their plant hosts (Gray and Gerdemann, 1973; Allen and Shachar-Hill, 2009), and that accumulation of plant sulphate transporter transcripts increases in colonized roots (Casieri et al., 2012; Giovannetti et al., 2014). Furthermore, the promoter of the *Lotus japonicus* sulphate transporter gene *LjSULTR1;2* is active in arbusculated cells (Giovannetti et al., 2014), suggesting a function in uptake from the peri-arbuscular space analogous to that of the PT4 high-affinity P transporter. With regard to Mg, transcripts encoding Mg transporters are known to accumulate to higher levels in wheat following mycorrhizal inoculation (Li et al., 2018). In common with P, the elements S, Ca and Mg are often poorly available due to low-mobility and formation of conjugates with other soil compounds (Kelly and Barber, 1991; Scherer, 2001; Lynch, 2019). As such, hyphal foraging and delivery of these nutrients by mycorrhizal fungi would be predicted to be of potential benefit to plants under field conditions. In the future, it will be informative to combine evaluation of RSA in the field with assessment of AM fungal communities and quantification of elements in reproductive stage plants and grain.

---

**Figure S1. Impact of AM colonization of growth, root architecture and nutrient levels.** The maize inbred lines B73 and W22 were grown with (M) or without (NC) inoculation with the AM fungus *Rhizophagus irregularis*. Traits are described in main text and Table S1, S2, S3. Boxes show 1st quartile, median and 3rd quartile. Whiskers extend to the most extreme points within 1.5x box length; outlying values beyond this range are not shown. Line segments show the reaction norm for each genotype, connecting median values for NC and M.

## ACKNOWLEDGEMENTS

We thank Greg Ziegler and Ivan Baxter (Danforth Center, MO) for ionomic analysis and Benjamin Barrales Gámez for assistance in the greenhouse experiment. We thank the reviewers for constructive feedback on the manuscript. M. Rosario Ramírez-Flores was supported by the Mexican National Council of Science and Technology (CONACYT) through its PhD scholarship program. This work was supported by the Mexican National Commission for the Study and Use of Biodiversity (CONABIO) project *Impact of native arbuscular mycorrhizal fungi on maize performance*.

## AUTHOR CONTRIBUTIONS

RRF, RJHS and VOP designed the study. MRRF conducted the experiments. MRRF, EBB and RRA performed image analysis. All authors contributed to data analysis and writing of the manuscript.

## PREPRINT

This article is available as a preprint at https://www.biorxiv.org/content/10.1101/695411v1

## CONFLICT OF INTEREST STATEMENT

Rubén Rellán-Álvarez is a review editor for Plant Direct.

