## Supplementary figures and images for "Inoculation with the mycorrhizal fungus *Rhizophagus irregularis* modulates the relationship between root growth and nutrient content in maize (*Zea mays* ssp. *mays* L.)"

### Supplemental Figure 1

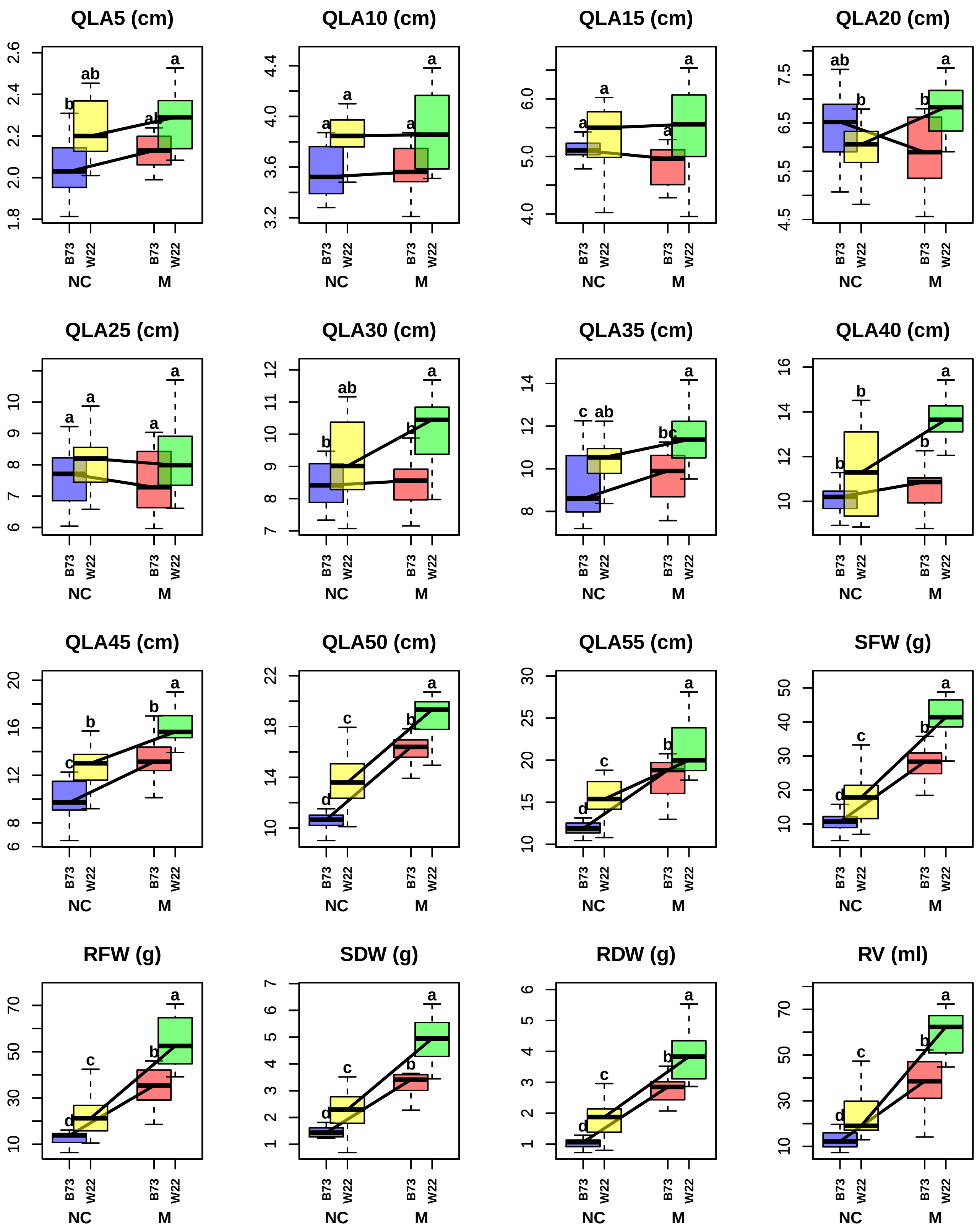

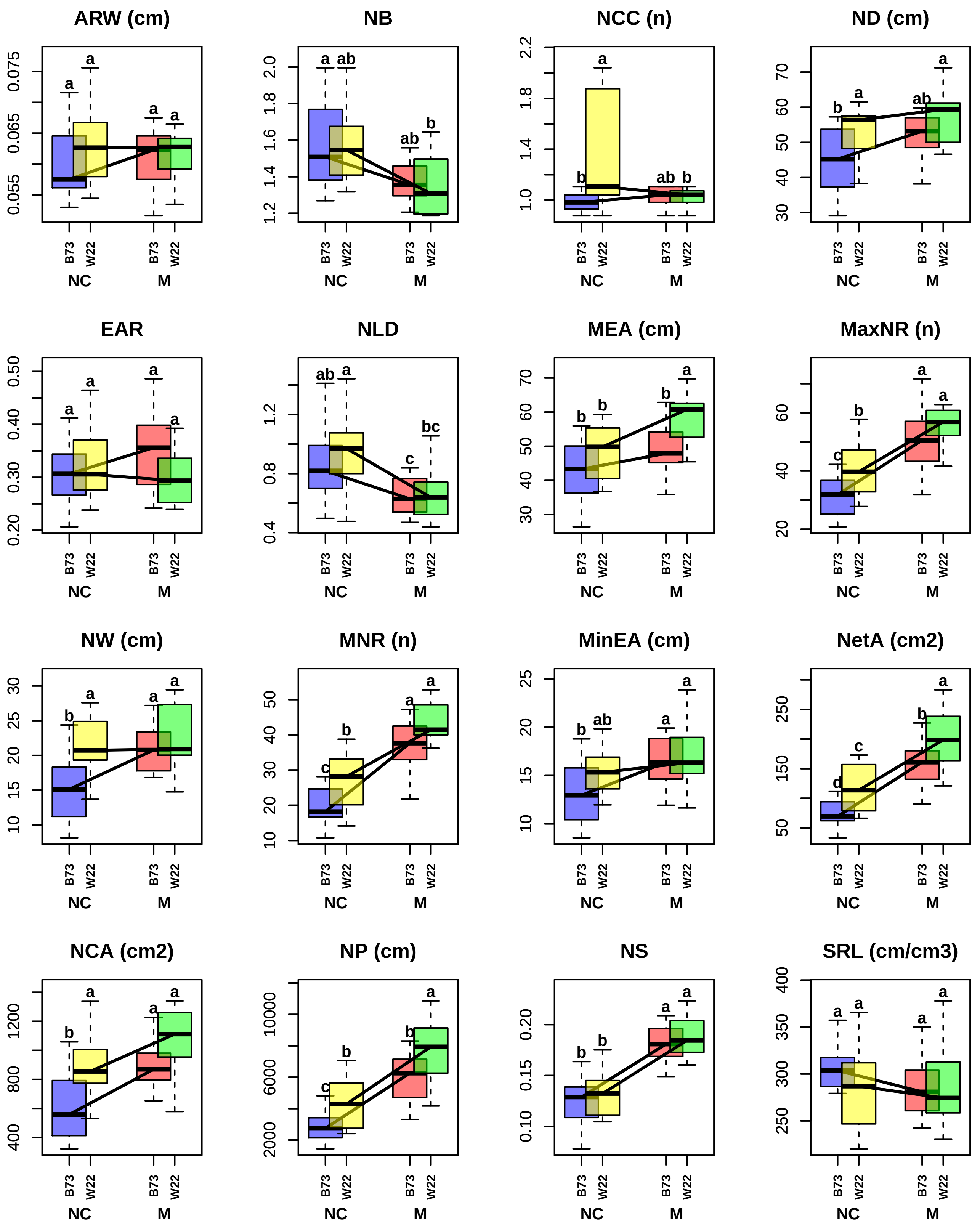

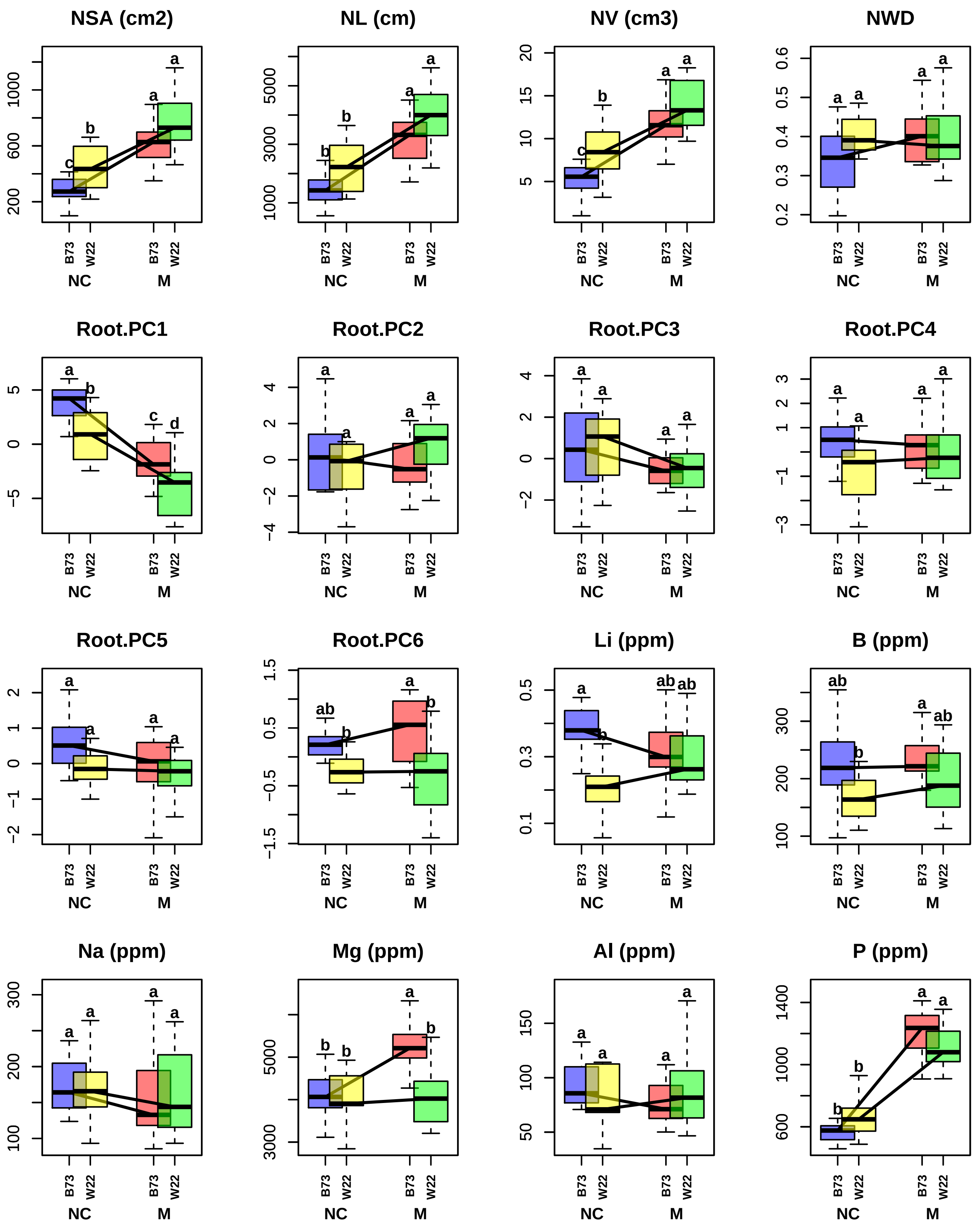

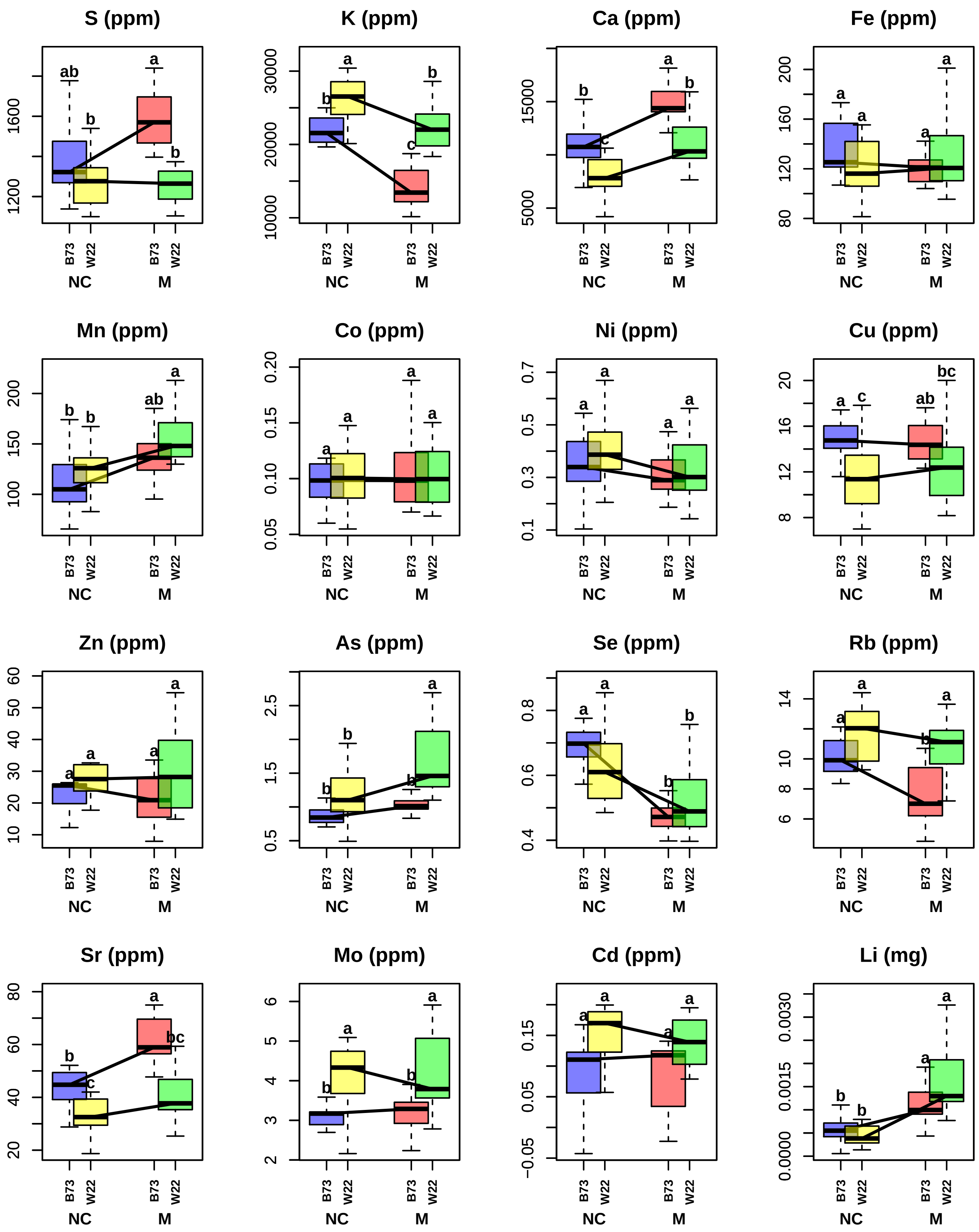

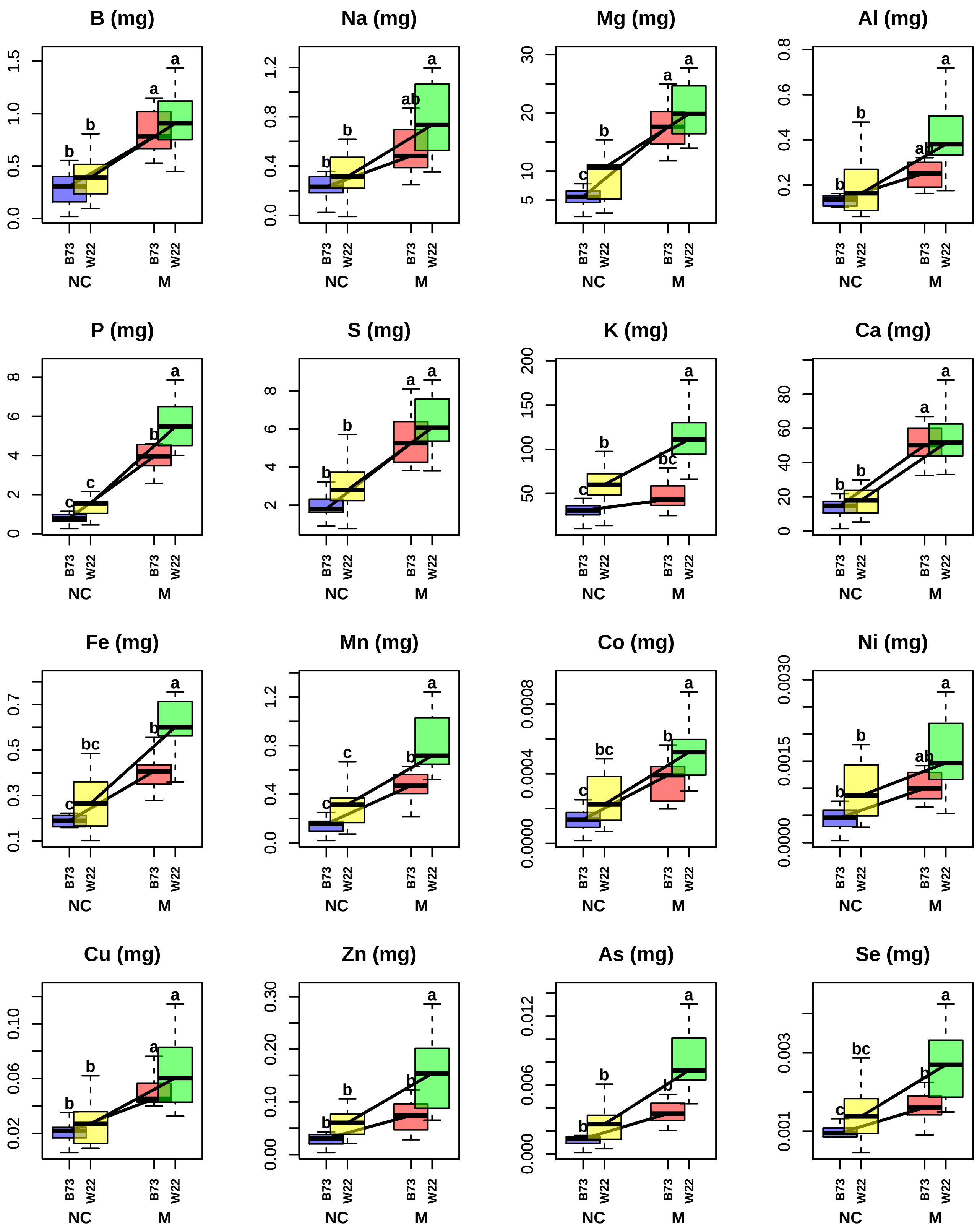

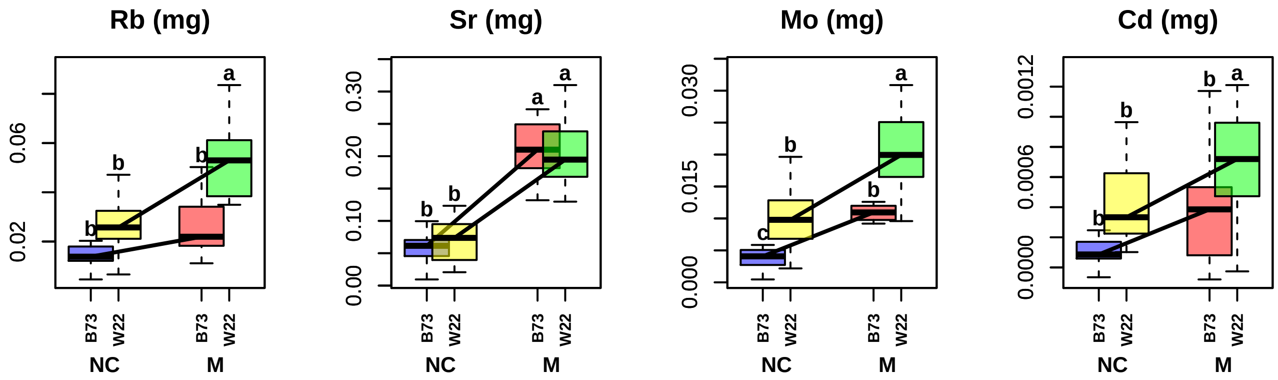
