## Supplemental Table 1 for "Inoculation with the mycorrhizal fungus *Rhizophagus irregularis* modulates the relationship between root growth and nutrient content in maize (*Zea mays* ssp. *mays* L.)"

**Table S1. GiA Roots traits (Galkovskyi et al. 2012).**

| Trait | Trait Description | Units |
|---|---|---|
| ARW | Average root width | cm |
| EAR | Ellipse axis ratio | NA |
| MEA | Major ellipse axis | cm |
| MaxNR | Maximum number of roots | n |
| MNR | Median number of roots | n |
| MinEA | Minor ellipse axis | cm |
| NetA | Network area | cm2 |
| NB | Network bushiness | NA |
| NCA | Network convex area | cm2 |
| NCC | Number of connected components | n |
| ND | Network depth | cm |
| NL | Network length | cm |
| NLD | Network length distribution | NA |
| NP | Network perimeter | cm |
| NS | Network solidity | NA |
| NSA | Network surface area | cm2 |
| NV | Network volume | cm3 |
| NW | Network width | cm |
| NWD | Network width to depth ratio | NA |
| SRL | Specific root length | cm/cm3 |
