## Supplemental Table 2 for "Inoculation with the mycorrhizal fungus *Rhizophagus irregularis* modulates the relationship between root growth and nutrient content in maize (*Zea mays* ssp. *mays* L.)"

| ID | Line | Block | Planting | Fungus | Hyphae | Vesicles | Arbuscules | QLA5 |
|---|---|---|---|---|---|---|---|---|
| MZ71002 | W22 | 1 | A | M | 80 | 65.2 | 77 | 2.349857143 |
| MZ71005 | W22 | 1 | A | M | 94.8 | 65.9 | 74.8 | 2.139857143 |
| MZ71007 | W22 | 1 | A | M | 95.6 | 63 | 57 | 2.209857143 |
| MZ71010 | W22 | 1 | A | M | 87.4 | 47.4 | 59.3 | 2.389857143 |
| MZ71038 | W22 | 4 | B | M | 92.6 | 63.7 | 67.4 | 2.493482143 |
| MZ71041 | W22 | 4 | B | M | 91.9 | 55.6 | 64.4 | 2.333482143 |
| MZ71043 | W22 | 4 | B | M | 85.2 | 41.5 | 59.3 | 2.113482143 |
| MZ71046 | W22 | 4 | B | M | 93.3 | 48.9 | 53.3 | 2.083482143 |
| MZ71055 | W22 | 5 | C | M | 91.1 | 54.8 | 68.1 | 2.138626374 |
| MZ71086 | W22 | 8 | D | M | 94.8 | 75.6 | 56.3 | 2.346190476 |
| MZ71089 | W22 | 8 | D | M | 89.6 | 51.9 | 63.7 | 2.526190476 |
| MZ71091 | W22 | 8 | D | M | 97.8 | 83.7 | 40.7 | 2.246190476 |
| MZ71014 | W22 | 2 | A | NC | 0 | 0 | 0 | 2.079857143 |
| MZ71017 | W22 | 2 | A | NC | 0 | 0 | 0 | 2.009857143 |
| MZ71019 | W22 | 2 | A | NC | 0 | 0 | 0 | 2.189857143 |
| MZ71022 | W22 | 2 | A | NC | 0 | 0 | 0 | 2.209857143 |
| MZ71026 | W22 | 3 | B | NC | 0 | 0 | 0 | 2.353482143 |
| MZ71029 | W22 | 3 | B | NC | 0 | 0 | 0 | 2.453482143 |
| MZ71031 | W22 | 3 | B | NC | 0 | 0 | 0 | 2.413482143 |
| MZ71034 | W22 | 3 | B | NC | 0 | 0 | 0 | 2.133482143 |
| MZ71062 | W22 | 6 | C | NC | 0 | 0 | 0 | 2.368626374 |
| MZ71065 | W22 | 6 | C | NC | 0 | 0 | 0 | 2.418626374 |
| MZ71067 | W22 | 6 | C | NC | 0 | 0 | 0 | 2.118626374 |
| MZ71077 | W22 | 7 | D | NC | 0 | 0 | 0 | 2.226190476 |
| MZ71079 | W22 | 7 | D | NC | 0 | 0 | 0 | 2.126190476 |
| MZ71082 | W22 | 7 | D | NC | 0 | 0 | 0 | 2.176190476 |

| QLA10 | QLA15 | QLA20 | QLA25 | QLA30 | QLA35 | QLA40 | QLA45 | QLA50 |
|---|---|---|---|---|---|---|---|---|
| 2.372071429 | 4.086309524 | 7.231642857 | 7.510428571 | 9.222607143 | 11.49228571 | 11.15433333 | 15.22372619 | 19.45614286 |
| 4.132071429 | 3.956309524 | 6.691642857 | 7.110428571 | 8.842607143 | 11.73228571 | 13.92433333 | 15.56372619 | 14.94614286 |
| 4.022071429 | 6.076309524 | 6.861642857 | 6.610428571 | 9.532607143 | 12.07228571 | 14.31433333 | 19.00372619 | 19.22614286 |
| 4.382071429 | 6.536309524 | 7.131642857 | 7.430428571 | 10.70260714 | 12.84228571 | 15.42433333 | 16.20372619 | 20.18614286 |
| 4.199821429 | 6.204017857 | 6.795892857 | 9.335178571 | 11.68348214 | 12.38803571 | 12.05625 | 15.68964286 | 19.43401786 |
| 3.709821429 | 5.584017857 | 5.905892857 | 8.155178571 | 7.973482143 | 10.76803571 | 13.67625 | 14.07964286 | 16.19401786 |
| 3.509821429 | 5.534017857 | 5.965892857 | 8.485178571 | 10.67348214 | 11.24803571 | 14.22625 | 17.78964286 | 20.71401786 |
| 3.789821429 | 5.654017857 | 5.975892857 | 8.445178571 | 10.15348214 | 10.87803571 | 13.45625 | 13.91964286 | 16.74401786 |
| 3.660686813 | 5.185796703 | 7.221950549 | 7.257967033 | 10.68276099 | 9.974285714 | 13.37961538 | 15.61300824 | 19.01137363 |
| 0.431904762 | 4.812142857 | 6.92047619 | 10.12142857 | 10.22327381 | 9.520952381 | 12.835 | 15.11172619 | 18.78714286 |
| 4.321904762 | 6.062142857 | 7.64047619 | 7.821428571 | 11.25327381 | 10.25095238 | 13.625 | 16.26172619 | 19.97714286 |
| 3.921904762 | 5.372142857 | 6.70047619 | 10.70142857 | 10.98327381 | 14.16095238 | 18.375 | 21.19172619 | 19.92714286 |
| 3.912071429 | 5.656309524 | 6.111642857 | 8.710428571 | 8.582607143 | 10.65228571 | 9.114333333 | 13.07372619 | 10.96614286 |
| 3.842071429 | 5.436309524 | 6.031642857 | 8.900428571 | 8.282607143 | 10.92228571 | 13.10433333 | 13.35372619 | 15.06614286 |
| 3.972071429 | 5.776309524 | 6.451642857 | 9.870428571 | 9.412607143 | 12.23228571 | 14.51433333 | 18.56372619 | 17.93614286 |
| 2.252071429 | 3.786309524 | 6.551642857 | 6.580428571 | 8.452607143 | 10.38228571 | 8.854333333 | 11.73372619 | 10.10614286 |
| 3.759821429 | 5.404017857 | 6.085892857 | 8.545178571 | 10.57348214 | 10.94803571 | 12.92625 | 11.50964286 | 13.19401786 |
| 4.099821429 | 6.024017857 | 6.325892857 | 8.555178571 | 11.16348214 | 11.65803571 | 10.88625 | 13.32964286 | 14.84401786 |
| 3.849821429 | 4.024017857 | 6.235892857 | 8.505178571 | 7.453482143 | 10.41803571 | 9.50625 | 12.60964286 | 14.79401786 |
| 3.649821429 | 5.504017857 | 5.815892857 | 7.985178571 | 10.37348214 | 10.93803571 | 13.56625 | 13.75964286 | 15.92401786 |
| 4.000686813 | 5.795796703 | 5.681950549 | 8.007967033 | 10.22276099 | 9.784285714 | 12.51961538 | 14.29300824 | 13.60137363 |
| 4.050686813 | 5.975796703 | 5.391950549 | 7.697967033 | 10.76276099 | 13.33428571 | 13.69961538 | 15.72300824 | 15.93137363 |
| 3.480686813 | 3.575796703 | 4.811950549 | 7.097967033 | 7.072760989 | 9.844285714 | 9.339615385 | 11.15300824 | 13.37137363 |
| 3.821904762 | 5.602142857 | 4.91047619 | 7.441428571 | 7.66327381 | 9.610952381 | 11.695 | 12.95172619 | 12.34714286 |
| 3.811904762 | 4.982142857 | 5.91047619 | 8.391428571 | 8.61327381 | 9.780952381 | 9.195 | 9.19172619 | 13.58714286 |
| 3.941904762 | 5.492142857 | 6.79047619 | 6.791428571 | 9.45327381 | 8.370952381 | 10.305 | 11.59172619 | 10.82714286 |

| QLA55 | SFW | RL | RV | RFW | SDW | RDW | ARW | NB |
|---|---|---|---|---|---|---|---|---|
| 19.82783333 | 40.43505952 | 91.20297619 | 62.30940476 | 48.25267857 | 4.753630952 | 2.94697619 | 0.061423824 | 1.493714404 |
| 19.23783333 | 35.33505952 | 94.20297619 | 62.30940476 | 42.64267857 | 3.763630952 | 2.86697619 | 0.063577887 | 1.243714404 |
| 24.39783333 | 48.77505952 | 108.2029762 | 72.30940476 | 67.83267857 | 5.223630952 | 4.98697619 | 0.062122998 | 1.193714404 |
| 24.94783333 | 45.40505952 | 109.2029762 | 67.30940476 | 58.67267857 | 5.013630952 | 3.72697619 | 0.062755611 | 1.532175942 |
| 22.735 | 36.68276786 | 111.2633929 | 49.77232143 | 46.91330357 | 3.973839286 | 2.984642857 | 0.062743561 | 1.500881167 |
| 18.285 | 28.52276786 | 76.76339286 | 44.77232143 | 39.14330357 | 3.443839286 | 3.384642857 | 0.066471908 | 1.64404356 |
| 23.315 | 47.53276786 | 104.7633929 | 64.77232143 | 61.87330357 | 6.233839286 | 4.164642857 | 0.064755909 | 1.198871711 |
| 20.145 | 41.15276786 | 112.2633929 | 44.77232143 | 42.16330357 | 4.583839286 | 3.244642857 | 0.05693937 | 1.478658945 |
| 18.3025 | 41.62839286 | 74.50041209 | 52.11607143 | 47.64037088 | 4.884656593 | 3.951950549 | 0.063150372 | 1.186174312 |
| 17.62416667 | 43.31255952 | 105.1029762 | 57.2077381 | 56.7010119 | 5.781964286 | 3.941309524 | 0.056839028 | 1.353169263 |
| 19.53416667 | 40.68255952 | 94.10297619 | 67.2077381 | 67.5210119 | 5.311964286 | 4.531309524 | 0.05345901 | 1.188334098 |
| 28.10416667 | 65.30255952 | 108.1029762 | 69.7077381 | 70.5710119 | 8.641964286 | 5.531309524 | 0.064789368 | 1.262693073 |
| 14.15783333 | 16.21505952 | 101.7029762 | 12.90940476 | 21.46267857 | 2.143630952 | 1.83697619 | 0.060667218 | 1.317243816 |
| 15.29783333 | 18.38505952 | 71.20297619 | 17.30940476 | 19.18267857 | 2.393630952 | 1.90697619 | 0.062037594 | 1.393714404 |
| 17.03783333 | 28.89505952 | 96.20297619 | 47.30940476 | 42.43267857 | 3.513630952 | 2.82697619 | 0.062471144 | 1.335177819 |
| 10.79783333 | 11.58505952 | 56.20297619 | 17.30940476 | 15.81267857 | 1.783630952 | 1.38697619 | 0.066713749 | 1.543714404 |
| 14.865 | 17.17276786 | 93.96339286 | 19.77232143 | 21.03330357 | 2.243839286 | 2.124642857 | 0.057953793 | 1.593876336 |
| 15.455 | 20.37276786 | 117.1633929 | 27.27232143 | 26.81330357 | 2.533839286 | 2.364642857 | 0.05442787 | 1.675717769 |
| 14.045 | 18.83276786 | 95.26339286 | 29.77232143 | 25.53330357 | 2.343839286 | 1.904642857 | 0.062830258 | 1.548658945 |
| 18.475 | 25.81276786 | 113.2633929 | 34.77232143 | 32.75330357 | 3.133839286 | 2.144642857 | 0.057045977 | 1.685801802 |
| 18.1325 | 21.37839286 | 111.5004121 | 27.11607143 | 23.48037088 | 2.774656593 | 1.851950549 | 0.0579362 | 1.408981329 |
| 18.8025 | 33.23839286 | 116.0004121 | 39.61607143 | 37.14037088 | 4.894656593 | 2.961950549 | 0.069147119 | 1.511362282 |
| 14.6925 | 15.31839286 | 66.50041209 | 17.11607143 | 16.28037088 | 1.854656593 | 1.601950549 | 0.075615666 | 1.449332207 |
| 17.45416667 | 10.15255952 | 77.10297619 | 16.7077381 | 10.5510119 | 0.871964286 | 0.871309524 | 0.06619515 | 1.78550009 |
| 15.91416667 | 9.522559524 | 103.1029762 | 18.2077381 | 11.4410119 | 1.341964286 | 1.181309524 | 0.067233966 | 1.996026406 |
| 11.19416667 | 6.952559524 | 86.10297619 | 14.7077381 | 10.6410119 | 0.691964286 | 0.801309524 | 0.064013579 | 1.567454977 |

| NCC | ND | EAR | NLD | MEA | MaxNR | NW | MNR | MinEA |
|---|---|---|---|---|---|---|---|---|
| 1.04047619 | 49.74721708 | 0.372232295 | 0.560637645 | 50.39717594 | 59.80595238 | 27.70073243 | 39.75952381 | 18.3347955 |
| 1.04047619 | 56.39801073 | 0.261103576 | 0.655551537 | 54.88024006 | 51.80595238 | 19.84358957 | 41.75952381 | 14.44494907 |
| 1.04047619 | 61.22340756 | 0.305825541 | 0.506753157 | 63.60849079 | 62.80595238 | 27.92295465 | 52.75952381 | 19.55793368 |
| 1.04047619 | 71.22340756 | 0.328199823 | 0.439293189 | 61.60152569 | 76.80595238 | 29.43089116 | 49.75952381 | 20.19201065 |
| 0.982142857 | 61.20656232 | 0.249750725 | 0.651363623 | 63.20024965 | 52.65178571 | 21.21712727 | 36.20535714 | 15.95038527 |
| 0.982142857 | 48.34941946 | 0.254311858 | 1.055352659 | 45.48053441 | 41.65178571 | 14.75680981 | 26.20535714 | 11.64511404 |
| 0.982142857 | 58.33354645 | 0.39267105 | 0.507141757 | 61.78674211 | 54.65178571 | 26.89966695 | 47.20535714 | 23.86343525 |
| 0.982142857 | 60.66687978 | 0.302937232 | 1.03515612 | 60.41514565 | 57.65178571 | 20.59807965 | 40.20535714 | 18.24915493 |
| 0.876373626 | 46.65126165 | 0.343811756 | 0.53784836 | 47.06245033 | 50.26236264 | 20.61922968 | 41.12362637 | 16.07988616 |
| 1.107142857 | 60.36243451 | 0.239283087 | 0.763756555 | 61.17339255 | 56.83928571 | 16.73592191 | 41.14285714 | 14.32620439 |
| 1.107142857 | 50.31481546 | 0.284429736 | 0.720266396 | 57.00007488 | 61.83928571 | 20.2279854 | 51.14285714 | 16.30489323 |
| 1.107142857 | 63.86317856 | 0.240198754 | 0.626051091 | 69.73805925 | 56.83928571 | 22.63497945 | 44.14285714 | 16.34483949 |
| 2.04047619 | 57.54086788 | 0.285256786 | 0.539416751 | 54.45112589 | 41.80595238 | 19.9705737 | 31.75952381 | 15.54137018 |
| 1.04047619 | 44.27102661 | 0.303499922 | 0.964618149 | 40.52862935 | 45.80595238 | 16.31978005 | 32.75952381 | 11.96178881 |
| 1.04047619 | 52.38213772 | 0.275801312 | 0.799807408 | 49.55707306 | 51.80595238 | 19.33565306 | 38.75952381 | 13.61915289 |
| 1.04047619 | 38.27102661 | 0.324606799 | 1.042827475 | 41.09173805 | 27.80595238 | 13.68485941 | 17.75952381 | 12.92517387 |
| 0.982142857 | 57.39703851 | 0.273215878 | 1.249882897 | 50.08472048 | 35.65178571 | 20.58220664 | 23.20535714 | 13.70690341 |
| 0.982142857 | 60.65100676 | 0.272681286 | 0.682636404 | 58.11920228 | 55.65178571 | 22.28061933 | 34.20535714 | 15.90452295 |
| 1.982142857 | 56.53989565 | 0.370556744 | 1.075661663 | 50.54551528 | 37.65178571 | 24.88379394 | 25.20535714 | 18.3740394 |
| 1.982142857 | 58.73037184 | 0.304285216 | 0.917266453 | 55.78615367 | 57.65178571 | 25.48696854 | 35.20535714 | 16.89866759 |
| 0.876373626 | 56.90126165 | 0.313009215 | 0.476015252 | 59.30082193 | 47.26236264 | 27.57235468 | 33.12362637 | 18.18939542 |
| 1.876373626 | 56.21376165 | 0.307282255 | 1.071797201 | 49.48810538 | 47.26236264 | 23.90047968 | 31.12362637 | 14.98336941 |
| 1.876373626 | 41.80751165 | 0.405644073 | 1.113004147 | 36.72832893 | 32.26236264 | 18.54110468 | 22.12362637 | 15.10831272 |
| 1.107142857 | 48.30067856 | 0.371304781 | 1.441709989 | 40.40015085 | 33.83928571 | 20.10372945 | 18.14285714 | 15.97226719 |
| 1.107142857 | 61.55067856 | 0.238208744 | 0.944742411 | 55.34660562 | 29.83928571 | 20.86935445 | 14.14285714 | 12.93330907 |
| 1.107142857 | 50.80687896 | 0.46458756 | 0.974350918 | 39.00276857 | 32.83928571 | 25.14862032 | 20.14285714 | 19.84043396 |

| NetA | NCA | NP | NS | SRL | NSA | NL | NV | NWD |
|---|---|---|---|---|---|---|---|---|
| 163.7509273 | 1053.498943 | 6464.660999 | 0.16037026 | 296.7106706 | 629.4316401 | 3376.648538 | 11.58602499 | 0.575704591 |
| 160.7985463 | 939.7907046 | 6048.184809 | 0.177335327 | 266.1606975 | 616.566728 | 3178.823141 | 11.79937442 | 0.351887722 |
| 238.018249 | 1312.537996 | 9501.883222 | 0.189128705 | 285.4029909 | 913.497335 | 4990.251712 | 16.39058982 | 0.461750834 |
| 270.9073899 | 1297.295491 | 10860.64513 | 0.218817171 | 282.6487709 | 1031.375501 | 5613.569172 | 18.25810406 | 0.413548689 |
| 200.147447 | 1138.305841 | 7507.910396 | 0.171443581 | 265.0755684 | 760.5978268 | 3846.309458 | 14.48300547 | 0.346104815 |
| 120.8980138 | 578.4606656 | 4167.386587 | 0.207674843 | 230.229735 | 464.9530619 | 2192.801522 | 9.698970227 | 0.300915547 |
| 238.6085203 | 1340.975027 | 8769.815158 | 0.174069719 | 250.3507816 | 896.1674262 | 4416.769776 | 17.17905325 | 0.46609224 |
| 188.0746326 | 1119.585004 | 7669.846904 | 0.162914518 | 328.3880871 | 723.6178381 | 4004.468189 | 12.77468978 | 0.338602249 |
| 163.2999826 | 815.3034008 | 5892.619348 | 0.196709713 | 266.206478 | 652.8096062 | 3223.041024 | 13.83034297 | 0.444175674 |
| 197.9555175 | 968.1167905 | 8199.641566 | 0.199982021 | 345.7704544 | 724.2281064 | 3991.82214 | 11.51983804 | 0.287349141 |
| 198.9257871 | 1104.397213 | 8715.578074 | 0.179759022 | 377.8861264 | 735.2915706 | 4326.282457 | 11.01747873 | 0.394795484 |
| 282.9995482 | 1225.308275 | 9566.154716 | 0.223291438 | 251.917433 | 1158.257058 | 5468.185979 | 25.95271368 | 0.356597703 |
| 136.2427388 | 959.0800722 | 5364.248301 | 0.145083012 | 311.9321261 | 521.556098 | 2777.108855 | 9.67046599 | 0.346468753 |
| 104.5271934 | 646.4616545 | 3851.29592 | 0.164987487 | 290.9727425 | 401.9208451 | 2012.34695 | 7.959260586 | 0.374498717 |
| 148.0104385 | 873.900556 | 5625.629253 | 0.175183075 | 282.9330094 | 569.2775539 | 2966.458061 | 10.77139981 | 0.371680163 |
| 75.29060985 | 530.9718586 | 2436.343539 | 0.139237185 | 234.971246 | 290.4912275 | 1264.537426 | 6.468980708 | 0.365728785 |
| 108.8269632 | 802.2204287 | 4227.497698 | 0.125423461 | 316.0663033 | 420.0305677 | 2223.658665 | 7.96111656 | 0.358354623 |
| 157.0685858 | 1050.539401 | 6612.513571 | 0.142600572 | 365.6451946 | 603.7669663 | 3458.118982 | 10.3994514 | 0.367767031 |
| 118.3852902 | 1005.516347 | 4351.481825 | 0.107655628 | 267.3174706 | 449.3489036 | 2223.658665 | 8.883327354 | 0.444019104 |
| 172.9055724 | 1359.688431 | 7055.450079 | 0.119943599 | 330.7380624 | 661.6570185 | 3641.753903 | 11.69345633 | 0.437565391 |
| 156.9642892 | 1340.154353 | 6177.338098 | 0.114218906 | 308.6014928 | 596.8622255 | 3202.728524 | 10.85096879 | 0.485316815 |
| 158.3656564 | 969.8279369 | 5331.603723 | 0.160339846 | 241.8852159 | 620.8383094 | 2840.541024 | 13.89914441 | 0.426752728 |
| 82.41497282 | 575.529475 | 2871.213098 | 0.143397936 | 220.1461131 | 300.5649928 | 1389.556649 | 6.537223065 | 0.446213034 |
| 66.2200071 | 773.1820548 | 2409.045336 | 0.105242389 | 248.7117482 | 247.5574422 | 1207.560979 | 5.163966353 | 0.406234383 |
| 78.7571165 | 836.1042965 | 2748.810961 | 0.110860131 | 246.7713676 | 300.4537845 | 1417.435979 | 6.312007158 | 0.342463303 |
| 67.3357141 | 790.202958 | 2720.562201 | 0.104541841 | 293.7451684 | 218.5502859 | 1130.838013 | 3.160387497 | 0.475863039 |

| Crown | X15cm | X30cm | X45cm | X60cm | X75cm | X90cm | Root.PC1 | Root.PC2 |
|---|---|---|---|---|---|---|---|---|
| 3.406047619 | 12.44765476 | 11.86377381 | 11.26066667 | 5.079464286 | 2.585 | 1.610071429 | -3.02 | -2.25 |
| 4.186047619 | 11.09765476 | 7.28377381 | 8.160666667 | 7.409464286 | 1.715 | 2.790071429 | -2.2 | 1.24 |
| 6.906047619 | 15.40765476 | 9.82377381 | 9.430666667 | 8.189464286 | 11.385 | 6.690071429 | -6.99 | -0.09 |
| 5.196047619 | 11.29765476 | 11.85377381 | 11.71066667 | 8.969464286 | 6.235 | 3.410071429 | -7.62 | 0.15 |
| 4.546339286 | 9.540446429 | 8.235982143 | 6.909375 | 5.793214286 | 4.73 | 7.157946429 | -3.46 | 1.54 |
| 5.446339286 | 13.23044643 | 7.905982143 | 5.959375 | 3.953214286 | 1.58 | 1.067946429 | 1.06 | 2.36 |
| 4.776339286 | 10.43044643 | 8.605982143 | 10.549375 | 10.78321429 | 8.53 | 8.197946429 | -6.17 | -1.71 |
| 4.816339286 | 9.940446429 | 6.905982143 | 5.859375 | 6.103214286 | 3.93 | 4.607946429 | -3.32 | 1.14 |
| 7.448406593 | 11.59924451 | 6.206799451 | 6.988461538 | 7.471002747 | 8.241153846 | -0.314697802 | -2.14 | -0.4 |
| 5.460714286 | 15.1964881 | 9.012440476 | 7.523333333 | 4.604464286 | 4.819166667 | 10.08440476 | -3.6 | 3.05 |
| 4.340714286 | 13.2664881 | 16.38244048 | 9.023333333 | 6.674464286 | 7.939166667 | 9.894404762 | -4.55 | 1.32 |
| 6.400714286 | 14.9664881 | 13.15244048 | 8.553333333 | 5.944464286 | 6.799166667 | 14.75440476 | -7.48 | 2.56 |
| 1.336047619 | 3.837654762 | 3.35377381 | 4.410666667 | 3.959464286 | 2.305 | 2.260071429 | -0.28 | 0.34 |
| 2.446047619 | 6.617654762 | 5.16377381 | 1.760666667 | 0.919464286 | 0.665 | 1.610071429 | 2.06 | 0.42 |
| 3.456047619 | 9.077654762 | 10.53377381 | 10.40066667 | 4.969464286 | 2.115 | 1.880071429 | -1.26 | 1 |
| 3.116047619 | 5.777654762 | 3.24377381 | 1.150666667 | 0.249464286 | 0.665 | 1.610071429 | 4.3 | -0.03 |
| 2.766339286 | 5.120446429 | 4.165982143 | 3.699375 | 2.213214286 | 1.81 | 1.257946429 | 1.44 | 0.87 |
| 3.566339286 | 4.650446429 | 5.125982143 | 5.369375 | 3.393214286 | 2.28 | 2.427946429 | -1.85 | 0.86 |
| 2.916339286 | 5.120446429 | 5.895982143 | 4.899375 | 3.213214286 | 2.03 | 1.457946429 | 0.37 | -1.71 |
| 3.076339286 | 6.000446429 | 8.315982143 | 6.909375 | 3.113214286 | 3 | 2.337946429 | -2.45 | -0.38 |
| 2.208406593 | 3.289244505 | 3.936799451 | 4.178461538 | 3.001002747 | 2.721153846 | 4.145302198 | -2.07 | -1.62 |
| 3.848406593 | 10.08924451 | 9.456799451 | 6.148461538 | 3.591002747 | 2.501153846 | 1.505302198 | -1.41 | -0.12 |
| 2.848406593 | 4.889244505 | 4.696799451 | 4.008461538 | 0.481002747 | -0.328846154 | -0.314697802 | 3.44 | -2.04 |
| 0.840714286 | 5.196488095 | 4.022440476 | 2.013333333 | 1.594464286 | -0.270833333 | -2.845595238 | 3.85 | -1.41 |
| 2.400714286 | 3.206488095 | 2.182440476 | 3.003333333 | 1.754464286 | 0.109166667 | -1.215595238 | 2.71 | 0.87 |
| 1.640714286 | 2.156488095 | 3.872440476 | 3.563333333 | 2.004464286 | -0.260833333 | -2.335595238 | 2.91 | -3.7 |

| Root.PC3 | Root.PC4 | Root.PC5 | Root.PC6 | Al.CON | As.CON | B.CON | Ca.CON | Cd.CON |
|---|---|---|---|---|---|---|---|---|
| -0.24 | 0.73 | -0.78 | -1.4 | 96.80212455 | 1.268048628 | 133.2783742 | 12978.7065 | 0.091706449 |
| -1.14 | -0.37 | 0.46 | -0.67 | 46.60820415 | 1.854389349 | 195.0170089 | 12550.75578 | 0.191703046 |
| -0.67 | -0.11 | -0.05 | -0.99 | 66.95410733 | 2.497317789 | 205.3750382 | 15920.22421 | 0.517235465 |
| 0.01 | -0.4 | -0.79 | 0.79 | 72.31161635 | 1.701539723 | 232.9930363 | 12660.85534 | 0.158295017 |
| 0.27 | -1.28 | 0.33 | -0.02 | 120.4167321 | 1.100630778 | 113.1344495 | 9770.822955 | 0.11415501 |
| -2.11 | -1.03 | -1.5 | -0.27 | 91.25807785 | 1.336642501 | 264.4138545 | 9603.36816 | -0.00754883 |
| -0.91 | -1.14 | -0.2 | 0.14 | 59.1527304 | 1.202825255 | 133.1960576 | 8301.566961 | 0.078841955 |
| 1.65 | 0.36 | -0.47 | 0.3 | 115.9186853 | 1.495774725 | 167.6714988 | 10446.15611 | 0.156631968 |
| -2.53 | 0.67 | -0.23 | -0.4 | 80.28718255 | 2.383676005 | 293.6968664 | 10522.42564 | 0.13812691 |
| 0.37 | 1.32 | 0.19 | -0.23 | 170.6261421 | 1.422304123 | 180.7222897 | 9755.849595 | 0.195004639 |
| 0.2 | 3.01 | -0.01 | -1.38 | 47.67598184 | 1.329462092 | 170.3595763 | 7650.953387 | 0.135756694 |
| -1.64 | -1.56 | -0.28 | -0.03 | 83.05790514 | 2.691004551 | 255.8496121 | 10210.2564 | 0.140055855 |
| 0.89 | -0.06 | 1.87 | 0.63 | 49.74947572 | 1.212879352 | 210.8762386 | 10228.37633 | 0.10512695 |
| -0.95 | 0.81 | -0.13 | -0.04 | 112.6250751 | 0.988645218 | 197.0387438 | 7004.257405 | 0.107204488 |
| -0.8 | 0.14 | 0 | -0.64 | 82.56631967 | 1.403981017 | 201.7034162 | 8163.867446 | 0.172620778 |
| -1.57 | -0.93 | -0.44 | -0.07 | 44.60215072 | 0.491410544 | 132.0280452 | 4192.68958 | 0.057068598 |
| 1.91 | -0.27 | -0.17 | -0.48 | 98.50769493 | 0.963741172 | 229.9159767 | 7046.397899 | 0.342035853 |
| 2.89 | 1.07 | 0.71 | 0.22 | 34.70980432 | 1.240062212 | 158.4218899 | 8443.772623 | 0.185460463 |
| 1.3 | -1.5 | -0.42 | -0.36 | 204.1820904 | 1.437728602 | 162.4560489 | 10138.44616 | 0.411397605 |
| 2.8 | 0.07 | -0.13 | -0.45 | 68.21560561 | 1.940212661 | 114.7979841 | 9551.483304 | 0.199498698 |
| 2.13 | -0.28 | 1.63 | -0.41 | 72.07709945 | 0.929823638 | 190.9179782 | 6912.731051 | 0.12270982 |
| -0.37 | -2.25 | -0.89 | -0.6 | 114.160374 | 1.910771419 | 164.8962325 | 10632.31632 | 0.131022713 |
| -2.26 | -1.99 | -1 | -0.1 | 68.63775511 | 1.428653909 | 134.6767057 | 7643.852674 | 0.175413845 |
| 0.82 | -1.76 | -1.53 | -0.17 | 69.64808893 | 0.526511641 | 110.2825252 | 7143.026284 | 0.126691598 |
| 1.85 | -3.08 | 0.22 | -0.37 | 68.13314656 | 0.947414264 | 161.4588591 | 7903.74363 | 0.167386294 |
| 1.24 | -0.56 | -0.34 | 0.26 | 112.668812 | 0.79954938 | 186.6347419 | 7724.018312 | 0.188611978 |

| Co.CON | Cu.CON | Fe.CON | K.CON | Li.CON | Mg.CON | Mn.CON | Mo.CON | Na.CON |
|---|---|---|---|---|---|---|---|---|
| 0.077947563 | 8.170951621 | 140.4727612 | 20825.41909 | 0.269056998 | 5467.317161 | 134.8004066 | 3.809948325 | 108.0882859 |
| 0.079772039 | 12.40220484 | 95.4937862 | 18494.85293 | 0.207034653 | 4266.095178 | 173.7157651 | 4.408060934 | 93.27978389 |
| 0.105426177 | 20.00702483 | 113.3819609 | 22052.75851 | 0.261953441 | 5098.007112 | 236.9412502 | 5.910097562 | 155.5978046 |
| 0.125298618 | 13.20612697 | 118.0353513 | 20405.73521 | 0.237386337 | 3806.246084 | 212.9759865 | 4.897351275 | 99.71742325 |
| 0.076372104 | 8.196534396 | 152.899963 | 30723.88127 | 0.451757111 | 4015.847762 | 130.9872105 | 3.549403988 | 262.4173828 |
| 0.137933902 | 10.99497055 | 123.2747539 | 19199.25349 | 0.223366449 | 4048.676809 | 168.3754999 | 2.782798938 | 157.3998984 |
| 0.066431813 | 8.872085787 | 107.6571745 | 28596.23087 | 0.187590038 | 3300.702227 | 158.5713244 | 3.710797263 | 132.2567211 |
| 0.123286999 | 11.01776965 | 164.4393089 | 23360.55809 | 0.262778297 | 3661.279864 | 145.0612286 | 3.579053951 | 124.8245268 |
| 0.112506598 | 16.10631468 | 113.1164225 | 18353.21084 | 0.66755401 | 4598.869352 | 139.695558 | 5.243126502 | 244.707923 |
| 0.15021997 | 15.07944003 | 201.0729542 | 23911.40858 | 0.490013736 | 4031.808998 | 129.9266093 | 3.767134956 | 188.1488646 |
| 0.093746954 | 12.35115617 | 107.342526 | 21984.53229 | 0.247571739 | 3259.517853 | 150.8989261 | 3.365979021 | 123.0713178 |
| 0.092540128 | 13.24275101 | 123.8632411 | 24377.53876 | 0.27341599 | 3206.155441 | 143.6477493 | 5.52494041 | 257.3439124 |
| 0.102056324 | 13.16874128 | 111.039672 | 22559.32357 | 0.18325852 | 4870.068236 | 148.9268407 | 4.743250247 | 102.5324367 |
| 0.108287054 | 9.220899478 | 143.3491077 | 26371.61531 | 0.271235931 | 4571.214852 | 129.6089327 | 3.929352378 | 146.7972208 |
| 0.086791881 | 10.20294839 | 105.9902221 | 27769.07936 | 0.217102625 | 3568.443448 | 131.864433 | 4.120366571 | 175.4810582 |
| 0.059620122 | 6.999392522 | 81.43023706 | 30015.39754 | 0.202094248 | 2842.304539 | 58.13411933 | 2.160098993 | 155.3660343 |
| 0.216158833 | 13.65815401 | 130.2941564 | 29940.04952 | 0.165112718 | 4927.642714 | 111.5309387 | 3.441969842 | 166.4361818 |
| 0.054834455 | 12.41156096 | 82.92390651 | 22604.64409 | 0.24191182 | 3860.0495 | 142.0833301 | 4.667606129 | 189.8751529 |
| 0.190483696 | 17.82818117 | 206.9323236 | 26717.5757 | 0.338567049 | 4560.695368 | 167.2207877 | 4.89103104 | 200.9806913 |
| 0.122404117 | 13.45758651 | 114.7625725 | 24277.97361 | 0.228789152 | 3917.667852 | 117.6739695 | 5.007819626 | 192.2874508 |
| 0.082558191 | 9.193089896 | 109.4064485 | 26105.60173 | 0.1188497 | 3891.327747 | 125.1356121 | 4.625967914 | 93.2040126 |
| 0.099278507 | 12.68097045 | 142.0162519 | 24091.6783 | 0.056599214 | 3138.383074 | 136.1697162 | 4.012261252 | -2.025333843 |
| 0.101849857 | 8.757504716 | 97.87115911 | 28558.04484 | 0.221632515 | 4269.21883 | 89.9063457 | 3.676926306 | 221.737853 |
| 0.07823383 | 10.30811893 | 117.443942 | 30410.97645 | 0.156486399 | 3876.504008 | 82.8559922 | 2.498968256 | 143.8800707 |
| 0.098964371 | 9.347206087 | 123.8728927 | 26704.4708 | 0.170050939 | 3876.473712 | 126.6972204 | 5.090953754 | 165.0653805 |
| 0.147554223 | 13.73872155 | 155.3487801 | 20118.02863 | 0.412398094 | 4021.023836 | 121.6859256 | 4.542862631 | 264.07288 |

| Ni.CON | P.CON | Rb.CON | S.CON | Se.CON | Sr.CON | Zn.CON | Al.TOT | As.TOT |
|---|---|---|---|---|---|---|---|---|
| 0.562681612 | 1005.447564 | 7.795813474 | 1251.293619 | 0.461932637 | 53.39864503 | 15.75942377 | 0.460161576 | 0.006027835 |
| 0.142708026 | 1112.902918 | 9.283447743 | 1300.821757 | 0.396716112 | 49.1355249 | 38.22085139 | 0.17541608 | 0.006979237 |
| 0.530793754 | 1031.19236 | 10.05092144 | 1589.214463 | 0.492516774 | 59.33037922 | 54.71846318 | 0.349743547 | 0.013045067 |
| 0.213646616 | 1046.852674 | 7.19929813 | 1123.409802 | 0.576890058 | 44.46893293 | 42.11171796 | 0.362543758 | 0.008530892 |
| 0.316743635 | 1007.321565 | 13.64187307 | 1273.335978 | 0.436024107 | 41.23721102 | 41.26274393 | 0.478516741 | 0.00437373 |
| 0.244473083 | 1226.966285 | 11.53740318 | 1103.59615 | 0.447746289 | 37.67673825 | 18.93456608 | 0.314278154 | 0.004603182 |
| 0.258450093 | 1203.214278 | 13.40107179 | 1373.516563 | 0.644352056 | 33.67099485 | 14.91034819 | 0.368748615 | 0.007498219 |
| 0.315402656 | 1355.914343 | 11.70840334 | 1261.735971 | 0.75683635 | 37.78220059 | 18.00491048 | 0.531352624 | 0.006856391 |
| 0.305327886 | 1136.438263 | 10.70469346 | 1267.256826 | 0.410182832 | 35.29861166 | 21.69769079 | 0.392175316 | 0.011643439 |
| 0.297341946 | 1042.442636 | 11.77779201 | 1352.379724 | 0.48715107 | 35.36927784 | 29.4615386 | 0.98655426 | 0.008223712 |
| 0.272226675 | 1276.832452 | 10.05800082 | 1254.21008 | 0.596570729 | 25.33089145 | 36.22565815 | 0.253253113 | 0.007062055 |
| 0.922306931 | 909.0701545 | 12.020592 | 845.3346936 | 0.490660147 | 35.82889431 | 26.92841165 | 0.71778345 | 0.023255565 |
| 0.669012601 | 720.0819157 | 9.852178285 | 1258.377355 | 0.854614345 | 42.00263838 | 23.7763816 | 0.106644516 | 0.002599966 |
| 0.205163951 | 654.9782187 | 13.56483383 | 1099.753053 | 0.485284619 | 31.60437844 | 24.16723393 | 0.269582866 | 0.002366452 |
| 0.221050227 | 611.2958837 | 13.16149168 | 1177.541578 | 0.816176508 | 33.53177495 | 30.07554616 | 0.290107576 | 0.004933071 |
| 0.370566541 | 811.391081 | 13.73768542 | 1419.747492 | 0.528879435 | 18.70155108 | 17.73886422 | 0.079553777 | 0.000876495 |
| 0.637326616 | 683.1132204 | 11.42683375 | 1294.569449 | 0.583038208 | 31.51681001 | 27.80523315 | 0.221035436 | 0.00216248 |
| 0.376063222 | 647.084085 | 10.19242106 | 1308.358958 | 0.572848392 | 36.56440136 | 21.08124022 | 0.087949066 | 0.003142118 |
| 0.621918628 | 487.7191044 | 11.7743182 | 1539.46251 | 0.625903044 | 40.55706808 | 32.62000159 | 0.478570005 | 0.003369805 |
| 0.470499335 | 648.7153809 | 9.272093059 | 1342.980593 | 0.594054828 | 39.36160189 | 32.11519867 | 0.213776745 | 0.006080315 |
| 0.397204598 | 572.1707513 | 12.66582234 | 1343.798678 | 0.639694351 | 25.88491007 | 24.62270642 | 0.199989199 | 0.002579941 |
| 0.368931697 | 523.1411044 | 9.625307784 | 1167.70556 | 0.693298359 | 41.59832232 | 32.06973035 | 0.558775827 | 0.00935257 |
| 0.33058789 | 513.4243311 | 12.38231259 | 1210.096964 | 0.526080002 | 33.4487744 | 20.61752794 | 0.127299465 | 0.002649662 |
| 0.472465476 | 929.3319774 | 14.40822356 | 1128.239148 | 0.526163281 | 29.36489909 | 27.26383358 | 0.060730646 | 0.000459099 |
| 0.247003214 | 768.8665485 | 12.30966725 | 1353.702073 | 0.698884686 | 29.45017073 | 51.62760246 | 0.091432249 | 0.001271396 |
| 0.409523869 | 647.711476 | 9.576162563 | 1131.421701 | 0.697599629 | 29.87322497 | 30.51924159 | 0.077962794 | 0.00055326 |

| B.TOT | Ca.TOT | Cd.TOT | Co.TOT | Cu.TOT | Fe.TOT | K.TOT | Li.TOT | Mg.TOT |
|---|---|---|---|---|---|---|---|---|
| 0.633556205 | 61.69598093 | 0.000435939 | 0.000370534 | 0.038841689 | 0.667755666 | 98.99635679 | 0.001278998 | 25.98960809 |
| 0.733972051 | 47.23641292 | 0.0007215 | 0.000300233 | 0.046677322 | 0.35940337 | 69.60780095 | 0.000779202 | 16.05600786 |
| 1.072803406 | 83.16137594 | 0.002701847 | 0.000550707 | 0.104509314 | 0.592265521 | 115.1954719 | 0.001368348 | 26.63010775 |
| 1.168141098 | 63.4768562 | 0.000793633 | 0.000628201 | 0.066210647 | 0.591785691 | 102.3068257 | 0.001190167 | 19.08311318 |
| 0.44957812 | 38.82768011 | 0.000453634 | 0.00030349 | 0.03257171 | 0.60759988 | 122.0917664 | 0.00179521 | 15.9583336 |
| 0.91059882 | 33.07245654 | -0.000026 | 0.000475022 | 0.037864912 | 0.424538441 | 66.11914343 | 0.000769238 | 13.94299225 |
| 0.830322816 | 51.75063425 | 0.000491488 | 0.000414125 | 0.055307157 | 0.671117524 | 178.2643074 | 0.001169406 | 20.57604721 |
| 0.768579203 | 47.88350077 | 0.000717976 | 0.000565128 | 0.050503685 | 0.753763364 | 107.0810439 | 0.001204533 | 16.78271848 |
| 1.434608335 | 51.39843579 | 0.000674703 | 0.000549556 | 0.078673816 | 0.552534879 | 89.64913234 | 0.003260772 | 22.4638975 |
| 1.044929825 | 56.40797394 | 0.00112751 | 0.000868566 | 0.087188784 | 1.16259664 | 138.2549104 | 0.002833242 | 23.31177564 |
| 0.904943985 | 40.64159115 | 0.000721135 | 0.00049798 | 0.0656089 | 0.570199664 | 116.7810503 | 0.001315092 | 17.31444243 |
| 2.21104321 | 88.23667119 | 0.001210358 | 0.000799728 | 0.114443381 | 1.070421706 | 210.6698193 | 0.002362851 | 27.70748082 |
| 0.452040832 | 21.92586409 | 0.000225353 | 0.000218771 | 0.028228921 | 0.238028078 | 48.35886427 | 0.000392839 | 10.43962901 |
| 0.471638036 | 16.76560732 | 0.000256608 | 0.000259199 | 0.02207143 | 0.343124861 | 63.12391467 | 0.000649239 | 10.94180136 |
| 0.708711366 | 28.68481735 | 0.000606526 | 0.000304955 | 0.035849395 | 0.372410525 | 97.57029676 | 0.000762819 | 12.53819335 |
| 0.235489308 | 7.478210909 | 0.000101789 | 0.00010634 | 0.012484333 | 0.145241491 | 53.5363921 | 0.000360462 | 5.069622353 |
| 0.515894501 | 15.81098443 | 0.000767473 | 0.000485026 | 0.030646703 | 0.292359147 | 67.18065933 | 0.000370486 | 11.05683831 |
| 0.401415608 | 21.39516279 | 0.000469927 | 0.000138942 | 0.031448901 | 0.210115852 | 57.27653524 | 0.000612966 | 9.780745068 |
| 0.38077087 | 23.76288841 | 0.00096425 | 0.000446463 | 0.041786391 | 0.48501611 | 62.62170355 | 0.000793547 | 10.68953697 |
| 0.359758432 | 29.93281361 | 0.000625197 | 0.000383595 | 0.042173913 | 0.359647458 | 76.08326748 | 0.000716988 | 12.27734142 |
| 0.529731827 | 19.18045479 | 0.000340478 | 0.000229071 | 0.025507667 | 0.303565324 | 72.43407997 | 0.000329767 | 10.79709819 |
| 0.807110432 | 52.04153719 | 0.000641311 | 0.000485934 | 0.062068996 | 0.695120784 | 117.920492 | 0.000277034 | 15.3613074 |
| 0.24977904 | 14.17672176 | 0.000325332 | 0.000188897 | 0.016242164 | 0.181517391 | 52.96536616 | 0.000411052 | 7.917934851 |
| 0.096162423 | 6.228463812 | 0.000110471 | 0.0000682 | 0.008988312 | 0.102406923 | 26.51728536 | 0.000136451 | 3.380173049 |
| 0.216672023 | 10.60654168 | 0.000224626 | 0.000132807 | 0.012543617 | 0.166232998 | 35.83644608 | 0.000228202 | 5.202089276 |
| 0.129144576 | 5.344744814 | 0.000130513 | 0.000102102 | 0.009506705 | 0.107495808 | 13.92095731 | 0.000285365 | 2.782404887 |

| Mn.TOT | Mo.TOT | Na.TOT | Ni.TOT | P.TOT | Rb.TOT | S.TOT | Se.TOT | Sr.TOT |
|---|---|---|---|---|---|---|---|---|
| 0.640791385 | 0.018111088 | 0.513811821 | 0.002674781 | 4.779526663 | 0.03705842 | 5.948188078 | 0.002195857 | 0.253837452 |
| 0.65380203 | 0.016590315 | 0.351070682 | 0.0005371 | 4.188555868 | 0.034939471 | 4.895813028 | 0.001493093 | 0.184927982 |
| 1.237693648 | 0.030872169 | 0.812785508 | 0.002772671 | 5.386568328 | 0.052502304 | 8.301469859 | 0.002572726 | 0.309920005 |
| 1.067782998 | 0.024553512 | 0.49994636 | 0.001071145 | 5.248532966 | 0.036094624 | 5.632362156 | 0.002892314 | 0.222950819 |
| 0.520522123 | 0.014104761 | 1.042804505 | 0.001258688 | 4.002934008 | 0.054210611 | 5.060032533 | 0.00173269 | 0.163870049 |
| 0.579858161 | 0.009583512 | 0.542059954 | 0.000841926 | 4.225474694 | 0.039732962 | 3.800607776 | 0.001541966 | 0.129752631 |
| 0.988508152 | 0.023132514 | 0.824467144 | 0.001611136 | 7.500644434 | 0.083540128 | 8.562281509 | 0.004016787 | 0.209899571 |
| 0.664937358 | 0.016405808 | 0.57217557 | 0.001445755 | 6.215293433 | 0.053669439 | 5.783594911 | 0.003469216 | 0.173187535 |
| 0.682364828 | 0.025610872 | 1.19531417 | 0.001491422 | 5.551110653 | 0.052288752 | 6.190114408 | 0.002003602 | 0.172421596 |
| 0.751231015 | 0.02178144 | 1.087870016 | 0.001719221 | 6.02736609 | 0.068098773 | 7.819411266 | 0.00281669 | 0.204503901 |
| 0.801569706 | 0.01787996 | 0.653750445 | 0.001446058 | 6.782488382 | 0.053427741 | 6.662319152 | 0.003168962 | 0.134556791 |
| 1.241398719 | 0.047746338 | 2.2239569 | 0.007970544 | 7.856151809 | 0.103881527 | 7.305352232 | 0.004240267 | 0.309632025 |
| 0.319244185 | 0.010167778 | 0.219791705 | 0.001434116 | 1.543589883 | 0.021119434 | 2.697496648 | 0.001831978 | 0.090038156 |
| 0.310235953 | 0.009405419 | 0.351378371 | 0.000491087 | 1.567776137 | 0.032469206 | 2.632402948 | 0.001161592 | 0.075649218 |
| 0.463322953 | 0.014477448 | 0.616575678 | 0.000776689 | 2.147868138 | 0.046244625 | 4.137446536 | 0.002867743 | 0.117818282 |
| 0.103689815 | 0.003852819 | 0.277115668 | 0.000660954 | 1.447222247 | 0.024502961 | 2.532305571 | 0.000943326 | 0.033356665 |
| 0.250257502 | 0.007723227 | 0.373456043 | 0.001430058 | 1.53279628 | 0.025639978 | 2.904805787 | 0.001308244 | 0.070718656 |
| 0.360016324 | 0.011826964 | 0.481113122 | 0.000952884 | 1.639607076 | 0.025825957 | 3.315171327 | 0.001451506 | 0.092648317 |
| 0.391938652 | 0.011463791 | 0.47106644 | 0.001457677 | 1.143135197 | 0.02759711 | 3.608252709 | 0.001467016 | 0.095059249 |
| 0.368771309 | 0.015693702 | 0.602597967 | 0.001474469 | 2.032969746 | 0.029057249 | 4.208685342 | 0.001861672 | 0.123352934 |
| 0.347208351 | 0.012835472 | 0.258609128 | 0.001102106 | 1.587577348 | 0.035143307 | 3.728579861 | 0.001774932 | 0.071821736 |
| 0.666503999 | 0.019638641 | -0.009913314 | 0.001805794 | 2.560596056 | 0.047112576 | 5.715517716 | 0.003393457 | 0.203609503 |
| 0.166745397 | 0.006819436 | 0.411247571 | 0.000613127 | 0.952225821 | 0.022964938 | 2.244314312 | 0.000975698 | 0.06203599 |
| 0.072247466 | 0.002179011 | 0.125458283 | 0.000411973 | 0.810344294 | 0.012563456 | 0.983784243 | 0.000458796 | 0.025605143 |
| 0.170023145 | 0.006831878 | 0.221511845 | 0.000331469 | 1.031791449 | 0.016519134 | 1.816619836 | 0.000937878 | 0.039521077 |
| 0.084202315 | 0.003143499 | 0.182729002 | 0.000283376 | 0.448193209 | 0.006626362 | 0.782903409 | 0.000482714 | 0.020671205 |

| Zn.TOT |
| ---: |
| 0.074914485 |
| 0.143849179 |
| 0.285829058 |
| 0.211132613 |
| 0.163971513 |
| 0.065207603 |
| 0.092948714 |
| 0.082531616 |
| 0.105985768 |
| 0.170345564 |
| 0.192429402 |
| 0.232714372 |
| 0.050967788 |
| 0.057847439 |
| 0.10567437 |
| 0.031639587 |
| 0.062390474 |
| 0.053416475 |
| 0.076456041 |
| 0.100643871 |
| 0.068319555 |
| 0.156970317 |
| 0.038238434 |
| 0.023773089 |
| 0.069282399 |
| 0.021118225 |

| ID | Line | Block | Planting | Fungus | Hyphae | Vesicles | Arbuscules | QLA5 |
|---|---|---|---|---|---|---|---|---|
| MZ71001 | B73 | 1 | A | M | 79.26 | 74.81 | 78.52 | 3.169857143 |
| MZ71004 | B73 | 1 | A | M | 69.63 | 52.59 | 47.41 | 1.989857143 |
| MZ71008 | B73 | 1 | A | M | 100 | 79.26 | 62.96 | 2.079857143 |
| MZ71037 | B73 | 4 | B | M | 97.78 | 82.22 | 51.85 | 2.043482143 |
| MZ71044 | B73 | 4 | B | M | 85.19 | 41.48 | 59.26 | 2.113482143 |
| MZ71047 | B73 | 4 | B | M | 97.78 | 83.7 | 70.37 | 2.083482143 |
| MZ71049 | B73 | 5 | C | M | 96.3 | 65.93 | 58.52 | 2.188626374 |
| MZ71052 | B73 | 5 | C | M | 95.56 | 61.48 | 71.85 | 2.238626374 |
| MZ71056 | B73 | 5 | C | M | 92.59 | 60.74 | 63.7 | 2.208626374 |
| MZ71085 | B73 | 8 | D | M | 89.63 | 48.89 | 69.63 | 1.836190476 |
| MZ71088 | B73 | 8 | D | M | 86.67 | 58.52 | 72.59 | 2.176190476 |
| MZ71095 | B73 | 8 | D | M | 97.04 | 61.48 | 82.22 | 2.146190476 |
| MZ71013 | B73 | 2 | A | NC | 0 | 0 | 0 | 1.819857143 |
| MZ71016 | B73 | 2 | A | NC | 0 | 0 | 0 | 2.029857143 |
| MZ71020 | B73 | 2 | A | NC | 2.22 | 0 | 0 | 2.079857143 |
| MZ71023 | B73 | 2 | A | NC | 0 | 0 | 0 | 1.919857143 |
| MZ71025 | B73 | 3 | B | NC | 0 | 0 | 0 | 1.813482143 |
| MZ71028 | B73 | 3 | B | NC | 0 | 0 | 0 | 2.013482143 |
| MZ71032 | B73 | 3 | B | NC | 0 | 0 | 0 | 2.153482143 |
| MZ71035 | B73 | 3 | B | NC | 0 | 0 | 0 | 2.133482143 |
| MZ71061 | B73 | 6 | C | NC | 0 | 0 | 0 | 1.988626374 |
| MZ71064 | B73 | 6 | C | NC | 0 | 0 | 0 | 2.188626374 |
| MZ71068 | B73 | 6 | C | NC | 0 | 0 | 0 | 2.308626374 |
| MZ71071 | B73 | 6 | C | NC | 0 | 0 | 0 | 1.888626374 |
| MZ71073 | B73 | 7 | D | NC | 0 | 0 | 0 | 2.246190476 |
| MZ71076 | B73 | 7 | D | NC | 0 | 0 | 0 | 1.986190476 |
| MZ71080 | B73 | 7 | D | NC | 0 | 0 | 0 | 2.096190476 |

| QLA10 | QLA15 | QLA20 | QLA25 | QLA30 | QLA35 | QLA40 | QLA45 | QLA50 |
|---|---|---|---|---|---|---|---|---|
| 3.262071429 | 4.436309524 | 6.651642857 | 6.830428571 | 8.172607143 | 10.45228571 | 9.704333333 | 13.07372619 | 16.36614286 |
| 3.822071429 | 5.096309524 | 5.411642857 | 7.820428571 | 7.152607143 | 7.572285714 | 9.724333333 | 10.11372619 | 12.57614286 |
| 3.792071429 | 5.196309524 | 5.631642857 | 8.250428571 | 9.882607143 | 8.992285714 | 10.94433333 | 14.57372619 | 17.82614286 |
| 3.459821429 | 3.364017857 | 5.385892857 | 7.185178571 | 8.863482143 | 8.708035714 | 11.52625 | 11.39964286 | 13.91401786 |
| 3.509821429 | 4.854017857 | 4.795892857 | 6.795178571 | 8.293482143 | 8.668035714 | 10.93625 | 14.33964286 | 17.19401786 |
| 3.209821429 | 4.584017857 | 6.795892857 | 9.035178571 | 7.493482143 | 10.60803571 | 10.48625 | 13.56964286 | 16.09401786 |
| 3.520686813 | 5.135796703 | 4.561950549 | 8.987967033 | 8.592760989 | 8.274285714 | 10.92961538 | 13.09300824 | 16.71137363 |
| 3.590686813 | 5.085796703 | 6.771950549 | 8.597967033 | 8.522760989 | 10.60428571 | 10.79961538 | 13.19300824 | 16.52137363 |
| 3.610686813 | 5.065796703 | 6.591950549 | 5.967967033 | 8.642760989 | 10.76428571 | 11.14961538 | 13.01300824 | 16.42137363 |
| 3.701904762 | 4.692142857 | 6.16047619 | 6.241428571 | 8.96327381 | 10.65095238 | 10.145 | 11.79172619 | 15.05714286 |
| 3.871904762 | 5.292142857 | 6.45047619 | 6.471428571 | 9.19327381 | 11.25095238 | 8.785 | 16.99172619 | 16.23714286 |
| 3.531904762 | 4.282142857 | 5.32047619 | 7.381428571 | 7.75327381 | 9.340952381 | 12.265 | 14.40172619 | 17.49714286 |
| 3.352071429 | 5.036309524 | 5.071642857 | 7.970428571 | 9.472607143 | 8.602285714 | 10.33433333 | 8.88372619 | 10.83614286 |
| 3.732071429 | 5.206309524 | 6.521642857 | 7.800428571 | 9.472607143 | 7.742285714 | 10.01433333 | 11.43372619 | 10.59614286 |
| 3.872071429 | 5.426309524 | 5.521642857 | 7.840428571 | 9.352607143 | 8.172285714 | 8.924333333 | 6.51372619 | 10.52614286 |
| 3.522071429 | 4.936309524 | 6.021642857 | 7.710428571 | 9.012607143 | 8.002285714 | 10.42433333 | 9.16372619 | 11.51614286 |
| 3.419821429 | 5.024017857 | 6.945892857 | 6.985178571 | 8.203482143 | 8.258035714 | 10.32625 | 11.99964286 | 10.68401786 |
| 3.279821429 | 4.784017857 | 5.785892857 | 6.835178571 | 7.513482143 | 7.358035714 | 9.72625 | 8.999642857 | 10.66401786 |
| 3.539821429 | 5.214017857 | 7.275892857 | 6.325178571 | 7.813482143 | 10.58803571 | 9.03625 | 12.26964286 | 10.15401786 |
| 3.459821429 | 5.104017857 | 7.185892857 | 6.875178571 | 8.273482143 | 10.63803571 | 9.62625 | 11.32964286 | 9.344017857 |
| 3.400686813 | 5.245796703 | 6.831950549 | 9.177967033 | 8.672760989 | 10.93428571 | 10.50961538 | 10.94300824 | 13.70137363 |
| 3.710686813 | 5.685796703 | 7.611950549 | 9.217967033 | 9.162760989 | 7.954285714 | 10.59961538 | 12.25300824 | 11.46137363 |
| 3.370686813 | 5.295796703 | 6.361950549 | 6.177967033 | 7.332760989 | 10.60428571 | 9.969615385 | 9.493008242 | 9.671373626 |
| 3.380686813 | 5.045796703 | 6.821950549 | 8.937967033 | 8.412760989 | 11.08428571 | 10.48961538 | 11.55300824 | 11.03137363 |
| 3.791904762 | 5.192142857 | 6.61047619 | 6.041428571 | 8.67327381 | 7.200952381 | 7.675 | 9.71172619 | 10.98714286 |
| 4.401904762 | 4.462142857 | 5.37047619 | 7.681428571 | 7.95327381 | 12.25095238 | 11.285 | 9.32172619 | 10.25714286 |
| 3.841904762 | 5.072142857 | 6.33047619 | 8.471428571 | 7.71327381 | 9.100952381 | 10.195 | 8.66172619 | 9.017142857 |

| QLA55 | SFW | RL | RV | RFW | SDW | RDW | ARW | NB |
|---|---|---|---|---|---|---|---|---|
| 16.56783333 | 24.16505952 | 98.20297619 | 27.30940476 | 26.98267857 | 2.993630952 | 2.43697619 | 0.067494371 | 1.303714404 |
| 15.83783333 | 18.41505952 | 89.20297619 | 19.80940476 | 18.57267857 | 2.273630952 | 1.54697619 | 0.057494371 | 1.453714404 |
| 18.24783333 | 28.08505952 | 103.2029762 | 37.30940476 | 38.52267857 | 3.293630952 | 2.88697619 | 0.057494371 | 1.463714404 |
| 16.255 | 21.48276786 | 84.26339286 | 34.77232143 | 31.22330357 | 2.473839286 | 2.074642857 | 0.061584669 | 1.548658945 |
| 15.825 | 27.26276786 | 68.26339286 | 34.77232143 | 32.96330357 | 3.513839286 | 2.884642857 | 0.061584669 | 1.288658945 |
| 19.555 | 28.48276786 | 78.76339286 | 39.77232143 | 33.02330357 | 3.423839286 | 2.434642857 | 0.051584669 | 1.558658945 |
| 20.3325 | 30.99839286 | 96.50041209 | 47.11607143 | 42.01037088 | 3.554656593 | 2.841950549 | 0.062963149 | 1.295647996 |
| 19.8925 | 30.77839286 | 92.50041209 | 47.11607143 | 42.20037088 | 3.644656593 | 3.151950549 | 0.062963149 | 1.335647996 |
| 19.4825 | 35.75839286 | 88.50041209 | 49.61607143 | 45.98037088 | 4.694656593 | 3.521950549 | 0.062963149 | 1.375647996 |
| 19.39416667 | 28.99255952 | 102.1029762 | 43.7077381 | 37.7110119 | 3.401964286 | 3.151309524 | 0.066142399 | 1.386026406 |
| 20.76416667 | 43.50255952 | 99.10297619 | 52.2077381 | 45.3710119 | 5.051964286 | 2.861309524 | 0.056142399 | 1.206026406 |
| 12.96416667 | 25.43255952 | 89.10297619 | 14.1077381 | 23.1410119 | 3.021964286 | 2.671309524 | 0.066142399 | 1.296026406 |
| 12.58783333 | 10.60505952 | 70.70297619 | 7.309404762 | 11.31267857 | 1.683630952 | 1.08697619 | 0.057494371 | 1.843714404 |
| 13.13783333 | 12.64505952 | 73.20297619 | 11.30940476 | 13.54267857 | 1.763630952 | 1.20697619 | 0.057494371 | 1.693714404 |
| 11.12783333 | 8.275059524 | 60.70297619 | 9.809404762 | 7.462678571 | 1.333630952 | 0.95697619 | 0.067494371 | 1.553714404 |
| 12.97783333 | 11.10505952 | 93.20297619 | 9.809404762 | 13.90267857 | 1.673630952 | 1.13697619 | 0.057494371 | 1.933714404 |
| 11.865 | 11.31276786 | 54.26339286 | 14.77232143 | 13.86330357 | 1.523839286 | 1.074642857 | 0.071584669 | 1.378658945 |
| 12.235 | 10.69276786 | 84.76339286 | 14.77232143 | 14.21330357 | 1.343839286 | 0.944642857 | 0.071584669 | 1.308658945 |
| 12.475 | 12.57276786 | 75.76339286 | 17.27232143 | 16.15330357 | 1.553839286 | 1.134642857 | 0.061584669 | 1.508658945 |
| 11.465 | 10.60276786 | 62.26339286 | 14.77232143 | 13.97330357 | 1.473839286 | 1.064642857 | 0.061584669 | 1.268658945 |
| 11.6725 | 11.81839286 | 83.50041209 | 19.61607143 | 15.41037088 | 1.394656593 | 1.261950549 | 0.052963149 | 1.495647996 |
| 12.4425 | 15.76839286 | 75.50041209 | 19.61607143 | 24.78037088 | 1.814656593 | 1.291950549 | 0.052963149 | 2.665647996 |
| 11.4025 | 9.648392857 | 55.50041209 | 12.11607143 | 11.36037088 | 1.224656593 | 0.901950549 | 0.062963149 | 1.305647996 |
| 11.2825 | 13.12839286 | 107.5004121 | 17.11607143 | 15.07037088 | 1.434656593 | 1.041950549 | 0.052963149 | 1.545647996 |
| 10.44416667 | 3.682559524 | 49.10297619 | 9.707738095 | 6.951011905 | 0.191964286 | 0.321309524 | 0.066142399 | 1.386026406 |
| 10.47416667 | 3.982559524 | 84.10297619 | 12.2077381 | 6.441011905 | 0.141964286 | 0.401309524 | 0.056142399 | 1.996026406 |
| 13.08416667 | 5.142559524 | 56.10297619 | 9.707738095 | 10.2310119 | 0.431964286 | 0.731309524 | 0.056142399 | 1.416026406 |

| NCC | ND | EAR | NLD | MEA | MaxNR | NW | MNR | MinEA |
|---|---|---|---|---|---|---|---|---|
| 1.04047619 | 52.9730901 | 0.38037497 | 0.801379774 | 47.32918078 | 43.80595238 | 21.7637483 | 33.75952381 | 17.53083125 |
| 1.04047619 | 47.0330901 | 0.38037497 | 0.741379774 | 45.14918078 | 31.80595238 | 22.8737483 | 21.75952381 | 16.58083125 |
| 1.04047619 | 57.7130901 | 0.36037497 | 0.571379774 | 53.47918078 | 62.80595238 | 22.3337483 | 42.75952381 | 18.83083125 |
| 0.982142857 | 51.67799089 | 0.25608553 | 0.778417466 | 46.07108565 | 40.65178571 | 17.38712727 | 27.20535714 | 11.92078793 |
| 0.982142857 | 38.17799089 | 0.42608553 | 0.518417466 | 35.84108565 | 52.65178571 | 16.81712727 | 42.20535714 | 14.56078793 |
| 0.982142857 | 45.83799089 | 0.48608553 | 0.838417466 | 40.90108565 | 71.65178571 | 24.54712727 | 47.20535714 | 19.02078793 |
| 0.876373626 | 56.37501165 | 0.296491611 | 0.536821285 | 56.67589219 | 49.26236264 | 23.90172968 | 37.12362637 | 16.15430998 |
| 1.876373626 | 55.51501165 | 0.276491611 | 0.756821285 | 52.32589219 | 49.26236264 | 18.18172968 | 36.12362637 | 14.27430998 |
| 0.876373626 | 53.37501165 | 0.416491611 | 0.586821285 | 45.18589219 | 53.26236264 | 19.79172968 | 38.12362637 | 18.79430998 |
| 1.107142857 | 59.83005356 | 0.351884668 | 0.53966227 | 54.92152662 | 60.83928571 | 27.18560445 | 43.14285714 | 19.91074122 |
| 1.107142857 | 58.38005356 | 0.241884668 | 0.46966227 | 62.83152662 | 51.83928571 | 19.21560445 | 42.14285714 | 14.70074122 |
| 1.107142857 | 50.04005356 | 0.311884668 | 0.66966227 | 48.48152662 | 42.83928571 | 16.90560445 | 32.14285714 | 15.48074122 |
| 1.04047619 | 38.7430901 | 0.22037497 | 1.111379774 | 38.95918078 | 20.80595238 | 8.143748299 | 10.75952381 | 8.550831248 |
| 1.04047619 | 55.8230901 | 0.26037497 | 0.981379774 | 52.78918078 | 30.80595238 | 14.7137483 | 17.75952381 | 13.70083125 |
| 1.04047619 | 35.4630901 | 0.33037497 | 0.921379774 | 38.29918078 | 24.80595238 | 12.0337483 | 15.75952381 | 12.04083125 |
| 1.04047619 | 47.2430901 | 0.41037497 | 1.411379774 | 43.87918078 | 31.80595238 | 20.7337483 | 15.75952381 | 17.27083125 |
| 0.982142857 | 35.88799089 | 0.28608553 | 0.818417466 | 32.66108565 | 22.65178571 | 10.05712727 | 17.20535714 | 9.230787929 |
| 0.982142857 | 57.26799089 | 0.34608553 | 0.718417466 | 54.70108565 | 22.65178571 | 21.46712727 | 18.20535714 | 18.79078793 |
| 0.982142857 | 46.18799089 | 0.27608553 | 0.678417466 | 47.35108565 | 36.65178571 | 17.54712727 | 25.20535714 | 12.95078793 |
| 0.982142857 | 39.94799089 | 0.26608553 | 0.668417466 | 41.09108565 | 25.65178571 | 12.16712727 | 21.20535714 | 11.01078793 |
| 0.876373626 | 52.07501165 | 0.356491611 | 0.676821285 | 45.64589219 | 42.26236264 | 24.38172968 | 28.12362637 | 16.28430998 |
| 0.876373626 | 55.38501165 | 0.206491611 | 1.976821285 | 55.04589219 | 39.26236264 | 10.40172968 | 16.12362637 | 10.76430998 |
| 0.876373626 | 39.37501165 | 0.266491611 | 0.926821285 | 34.37589219 | 32.26236264 | 10.31172968 | 24.12362637 | 9.17430998 |
| 0.876373626 | 57.24501165 | 0.306491611 | 0.496821285 | 55.96589219 | 40.26236264 | 18.70172968 | 26.12362637 | 16.58430998 |
| 1.107142857 | 29.11005356 | 0.341884668 | 0.74966227 | 26.41152662 | 31.83928571 | 15.11560445 | 22.14285714 | 10.09074122 |
| 1.107142857 | 45.24005356 | 0.341884668 | 0.99966227 | 43.31152662 | 35.83928571 | 17.91560445 | 17.14285714 | 15.29074122 |
| 1.107142857 | 35.04005356 | 0.411884668 | 0.73966227 | 30.18152662 | 36.83928571 | 15.38560445 | 25.14285714 | 13.67074122 |

| NetA | NCA | NP | NS | SRL | NSA | NL | NV | NWD |
|---|---|---|---|---|---|---|---|---|
| 143.0196776 | 967.4090493 | 5262.374809 | 0.14863586 | 251.5594434 | 542.7704188 | 2654.268855 | 10.58434362 | 0.415783983 |
| 90.36967763 | 762.3190493 | 3309.934809 | 0.11863586 | 301.9194434 | 349.9604188 | 1714.918855 | 7.014343623 | 0.505783983 |
| 186.7496776 | 1127.509049 | 7590.134809 | 0.16863586 | 312.3994434 | 717.4204188 | 3964.588855 | 12.75434362 | 0.385783983 |
| 125.59655 | 652.5600231 | 4442.849285 | 0.187946692 | 242.2392138 | 480.4575304 | 2309.432633 | 9.80764697 | 0.333806526 |
| 127.24655 | 511.9500231 | 4678.319285 | 0.257946692 | 269.0092138 | 494.6875304 | 2488.832633 | 9.73764697 | 0.443806526 |
| 160.93655 | 830.0900231 | 6696.629285 | 0.187946692 | 349.9792138 | 626.3075304 | 3537.752633 | 10.94764697 | 0.543806526 |
| 171.3562521 | 993.9261596 | 6453.968098 | 0.173670734 | 295.8729349 | 655.8007654 | 3368.398524 | 12.3403006 | 0.427240934 |
| 160.8462521 | 886.7261596 | 6271.968098 | 0.173670734 | 305.8029349 | 627.8507654 | 3326.668524 | 11.5403006 | 0.327240934 |
| 173.4862521 | 852.2561596 | 6235.748098 | 0.203670734 | 273.0029349 | 679.5207654 | 3317.888524 | 13.7503006 | 0.377240934 |
| 227.0632298 | 1226.941151 | 8304.922836 | 0.188799625 | 278.1493979 | 896.3552735 | 4511.827854 | 18.36154886 | 0.445683642 |
| 207.8032298 | 964.7811515 | 7675.422836 | 0.208799625 | 283.7193979 | 835.2952735 | 4283.687854 | 16.86154886 | 0.335683642 |
| 136.7032298 | 827.3711515 | 4710.002836 | 0.168799625 | 252.5393979 | 539.0852735 | 2555.967854 | 11.59154886 | 0.335683642 |
| 57.25967763 | 360.3390493 | 1843.804809 | 0.15863586 | 295.6094434 | 224.9204188 | 987.2188547 | 5.094343623 | 0.205783983 |
| 91.96967763 | 756.3590493 | 3491.504809 | 0.11863586 | 336.1194434 | 355.8504188 | 1829.408855 | 6.934343623 | 0.255783983 |
| 65.66967763 | 451.8690493 | 2157.234809 | 0.13863586 | 279.2394434 | 253.3404188 | 1118.598855 | 5.584343623 | 0.345783983 |
| 80.31967763 | 799.7690493 | 2873.944809 | 0.09863586 | 304.1194434 | 310.8204188 | 1505.618855 | 6.424343623 | 0.445783983 |
| 61.52655003 | 367.3600231 | 1937.989285 | 0.157946692 | 213.6592138 | 241.9975304 | 1066.022633 | 5.52764697 | 0.273806526 |
| 96.27655003 | 1058.610023 | 3315.399285 | 0.077946692 | 237.8792138 | 364.7875304 | 1691.072633 | 7.59764697 | 0.373806526 |
| 97.37655003 | 728.3200231 | 3804.069285 | 0.117946692 | 327.8092138 | 379.1575304 | 2021.582633 | 7.20764697 | 0.383806526 |
| 69.37655003 | 454.5800231 | 2572.209285 | 0.137946692 | 314.6392138 | 273.0675304 | 1394.812633 | 5.55764697 | 0.293806526 |
| 111.2462521 | 953.1961596 | 4602.198098 | 0.113670734 | 317.3929349 | 413.7107654 | 2322.168524 | 6.800300603 | 0.467240934 |
| 76.17625212 | 558.0861596 | 3346.918098 | 0.133670734 | 317.7929349 | 289.7307654 | 1733.778524 | 4.400300603 | 0.197240934 |
| 62.61625212 | 376.7861596 | 2707.038098 | 0.163670734 | 288.5029349 | 232.4207654 | 1351.678524 | 3.560300603 | 0.267240934 |
| 108.9162521 | 998.1061596 | 4813.558098 | 0.103670734 | 357.2129349 | 407.1607654 | 2449.008524 | 5.970300603 | 0.327240934 |
| 33.2032298 | 321.3211515 | 1436.542836 | 0.138799625 | 285.1493979 | 99.56527347 | 557.9478538 | 1.001548858 | 0.475683642 |
| 63.9832298 | 785.1811515 | 2741.482836 | 0.098799625 | 303.5493979 | 244.4952735 | 1425.737854 | 4.041548858 | 0.385683642 |
| 49.3732298 | 448.3811515 | 2117.292836 | 0.128799625 | 298.2493979 | 184.2252735 | 1086.197854 | 2.851548858 | 0.415683642 |

| Crown | X15cm | X30cm | X45cm | X60cm | X75cm | X90cm | Root.PC1 | Root.PC2 |
|---|---|---|---|---|---|---|---|---|
| 4.306047619 | 6.577654762 | 4.88377381 | 4.050666667 | 3.879464286 | 1.445 | 1.840071429 | -0.57 | -1.41 |
| 4.346047619 | 3.877654762 | 3.10377381 | 2.580666667 | 1.799464286 | 1.055 | 1.810071429 | 1.82 | -2.24 |
| 4.716047619 | 6.377654762 | 6.58377381 | 8.500666667 | 5.929464286 | 3.475 | 2.940071429 | -3.32 | -0.38 |
| 3.206339286 | 7.630446429 | 6.075982143 | 6.559375 | 4.423214286 | 2.1 | 1.227946429 | 0.91 | 1.57 |
| 4.036339286 | 8.820446429 | 7.025982143 | 8.659375 | 2.533214286 | 0.82 | 1.067946429 | 0.06 | -0.99 |
| 3.356339286 | 9.090446429 | 8.635982143 | 4.659375 | 3.913214286 | 2.3 | 1.067946429 | -2.11 | -2.75 |
| 3.958406593 | 10.15924451 | 8.766799451 | 7.748461538 | 3.021002747 | 3.961153846 | 4.395302198 | -2.56 | -0.02 |
| 4.098406593 | 11.69924451 | 5.566799451 | 5.898461538 | 6.131002747 | 5.001153846 | 3.805302198 | -1.61 | 1.49 |
| 4.958406593 | 10.11924451 | 5.426799451 | 7.978461538 | 11.70100275 | 4.881153846 | 0.915302198 | -2.4 | -0.66 |
| 3.450714286 | 7.196488095 | 0.042440476 | 6.623333333 | 7.724464286 | 6.799166667 | 5.874404762 | -4.82 | -1.05 |
| 4.690714286 | 9.456488095 | 6.482440476 | 8.813333333 | 5.604464286 | 4.239166667 | 6.084404762 | -3.85 | 2.16 |
| 5.420714286 | 3.926488095 | 5.072440476 | 4.343333333 | 5.844464286 | 1.629166667 | -3.095595238 | 0.23 | 0.3 |
| 2.276047619 | 4.597654762 | 1.20377381 | 0.440666667 | 0.519464286 | 0.665 | 1.610071429 | 5.87 | 3.23 |
| 1.756047619 | 3.607654762 | 2.62377381 | 2.020666667 | 1.259464286 | 0.665 | 1.610071429 | 2.94 | 1.78 |
| 1.256047619 | 1.157654762 | 1.71377381 | 0.770666667 | 0.289464286 | 0.665 | 1.610071429 | 5.09 | 0.03 |
| 2.306047619 | 4.027654762 | 3.15377381 | 1.050666667 | 0.719464286 | 1.035 | 1.610071429 | 3.18 | -1.77 |
| 2.346339286 | 4.250446429 | 3.115982143 | 1.629375 | 0.633214286 | 0.82 | 1.067946429 | 5.45 | 1.11 |
| 1.306339286 | 2.850446429 | 2.265982143 | 2.499375 | 2.513214286 | 1.71 | 1.067946429 | 1.92 | -1.76 |
| 1.966339286 | 4.120446429 | 3.315982143 | 2.519375 | 1.793214286 | 1.37 | 1.067946429 | 2.33 | 0.18 |
| 1.696339286 | 4.290446429 | 2.765982143 | 2.229375 | 1.103214286 | 0.82 | 1.067946429 | 4.22 | 1.16 |
| 1.988406593 | 3.319244505 | 3.726799451 | 4.008461538 | 1.751002747 | 0.641153846 | -0.024697802 | 0.89 | -1.75 |
| 2.828406593 | 8.739244505 | 10.90679945 | 2.388461538 | -0.258997253 | 0.221153846 | -0.044697802 | 4.29 | 4.47 |
| 2.268406593 | 2.949244505 | 3.776799451 | 3.148461538 | 0.191002747 | -0.658846154 | -0.314697802 | 4.92 | 1.66 |
| 1.918406593 | 2.989244505 | 2.786799451 | 3.118461538 | 1.651002747 | 1.511153846 | 1.095302198 | 0.7 | 0.13 |
| 2.210714286 | 3.376488095 | 2.782440476 | 2.963333333 | -0.075535714 | -1.210833333 | -3.095595238 | 6.03 | -1.58 |
| 1.550714286 | 2.346488095 | 2.342440476 | 2.383333333 | 0.954464286 | -0.040833333 | -3.095595238 | 3.88 | -0.74 |
| 2.400714286 | 4.336488095 | 3.762440476 | 3.353333333 | 0.684464286 | -1.210833333 | -3.095595238 | 4.69 | -1.77 |

| Root.PC3 | Root.PC4 | Root.PC5 | Root.PC6 | Al.CON | As.CON | B.CON | Ca.CON | Cd.CON |
|---|---|---|---|---|---|---|---|---|
| -0.71 | -1.29 | 0.12 | 0.4 | 107.1360494 | 0.954050675 | 179.7151302 | 15126.51553 | 0.391270476 |
| 0.94 | 0.47 | 0.48 | -0.46 | 78.64604942 | 0.904050675 | 265.7551302 | 14250.18553 | 0.041270476 |
| 0.75 | 0.88 | 0.01 | 0.97 | 76.14604942 | 0.984050675 | 221.5651302 | 17678.60553 | 0.051270476 |
| -1.11 | -0.53 | -0.17 | 0.08 | 65.73617832 | 1.15333994 | 213.5713051 | 15948.98223 | 0.027338125 |
| -3.54 | 2.21 | -1.06 | 1.06 | 72.31617832 | 1.00333994 | 229.3813051 | 12078.08223 | -0.022661875 |
| 0.43 | 3.61 | -2.09 | 0.66 | 50.11617832 | 1.02333994 | 221.7113051 | 14342.21223 | -0.002661875 |
| -0.35 | -0.08 | 0.71 | -0.53 | 69.90263751 | 0.832445137 | 249.3492364 | 14425.35912 | 0.140521568 |
| -0.53 | 0.19 | 0.29 | -0.24 | 69.58263751 | 1.012445137 | 315.1492364 | 14239.67912 | 0.120521568 |
| -1.64 | 0.37 | -0.85 | 1.16 | 111.9226375 | 1.012445137 | 212.9592364 | 13861.83912 | 0.120521568 |
| -0.46 | -0.81 | -0.09 | 0.8 | 59.46217651 | 1.257834504 | 216.8685077 | 18148.8019 | 0.124729373 |
| -0.65 | 0.52 | 1.04 | 0.45 | 55.49217651 | 1.027834504 | 206.3285077 | 13251.9819 | 0.124729373 |
| -1.29 | -0.88 | 0.75 | 0.96 | 205.2221765 | 1.507834504 | 343.2285077 | 15968.0519 | 0.114729373 |
| 0.12 | 0.5 | -0.12 | 0.53 | 95.03604942 | 0.774050675 | 254.2451302 | 8115.675527 | 0.051270476 |
| 2.02 | 0.16 | 0.8 | 0.56 | 77.42604942 | 0.854050675 | 218.7651302 | 10744.92553 | 0.041270476 |
| -1.27 | -0.2 | 0.2 | 0.24 | 361.4660494 | 0.704050675 | 204.0751302 | 6935.525527 | 0.061270476 |
| 2.61 | -0.34 | -1.64 | 0.15 | 83.31604942 | 0.774050675 | 193.9551302 | 8991.845527 | 0.051270476 |
| -3.29 | -1.21 | 0.51 | 0.25 | 84.76617832 | 0.84333994 | 354.6413051 | 10719.14223 | -0.042661875 |
| 0.43 | -3.25 | 1.87 | 0.15 | 253.1861783 | 1.02333994 | 184.2313051 | 11006.40223 | 0.457338125 |
| 0.6 | 0.63 | 1.25 | -0.44 | 70.81617832 | 1.03333994 | 261.4613051 | 13807.83223 | 0.077338125 |
| -0.96 | 1.03 | 1.75 | -0.05 | 98.75617832 | 0.88333994 | 209.0813051 | 9665.522231 | 0.167338125 |
| 1.79 | 1.2 | 0.72 | -0.11 | 74.01263751 | 1.132445137 | 283.7992364 | 12205.64912 | 0.130521568 |
| 3.85 | -0.21 | -3.57 | 0.12 | 76.41263751 | 0.832445137 | 304.4792364 | 9848.929122 | 0.110521568 |
| -1.74 | 1.01 | 0.72 | 0.21 | 132.7126375 | 0.762445137 | 97.12923643 | 9840.269122 | 0.110521568 |
| 3.07 | 1.03 | 2.08 | 0.67 | 85.68263751 | 1.082445137 | 139.8192364 | 15211.13912 | 0.110521568 |
| -2.14 | 1.33 | 0.39 | -0.97 | 121.2821765 | 0.767834504 | 99.42850768 | 11790.7019 | 0.134729373 |
| 2.38 | 0.26 | -0.48 | 0.29 | 96.37217651 | 0.887834504 | 239.4685077 | 10777.8819 | 0.114729373 |
| -0.7 | 2.22 | 0.14 | 0.41 | 71.47217651 | 0.727834504 | 266.4785077 | 12109.1419 | 0.104729373 |

| Co.CON | Cu.CON | Fe.CON | K.CON | Li.CON | Mg.CON | Mn.CON | Mo.CON | Na.CON |
|---|---|---|---|---|---|---|---|---|
| 0.188052939 | 14.55129945 | 142.2445425 | 14411.71836 | 0.337748332 | 5590.071559 | 140.4426753 | 3.070141536 | 120.2817707 |
| 0.098052939 | 17.56129945 | 122.4945425 | 12959.15836 | 0.287748332 | 5178.101559 | 95.30267529 | 2.740141536 | 108.8017707 |
| 0.098052939 | 13.31129945 | 129.5345425 | 12689.45836 | 0.297748332 | 5736.901559 | 129.9526753 | 3.260141536 | 133.1917707 |
| 0.080054474 | 16.63412942 | 119.7282162 | 10148.93109 | 0.624130732 | 5479.835075 | 138.7855753 | 3.905625507 | 291.543075 |
| 0.070054474 | 12.32412942 | 109.5482162 | 16137.19109 | 0.274130732 | 4040.525075 | 111.3355753 | 3.475625507 | 149.123075 |
| 0.070054474 | 15.48412942 | 104.0982162 | 10309.41109 | 0.264130732 | 5005.345075 | 153.6555753 | 3.255625507 | 222.783075 |
| 0.118333903 | 14.18610826 | 124.6953261 | 17043.30769 | 0.32909072 | 5244.129928 | 147.9320903 | 2.776639972 | 166.3503987 |
| 0.128333903 | 12.78610826 | 107.4353261 | 11857.75769 | 0.11909072 | 4956.739928 | 131.5520903 | 3.456639972 | 115.9203987 |
| 0.098333903 | 12.95610826 | 134.4053261 | 16791.08769 | 0.40909072 | 5314.959928 | 133.9020903 | 3.316639972 | 85.6103987 |
| 0.118332801 | 17.61041918 | 123.7675969 | 13907.02143 | 0.500791996 | 6325.836362 | 185.1844577 | 3.455129101 | 255.426588 |
| 0.078332801 | 15.11041918 | 109.8275969 | 18745.12143 | 0.240791996 | 4271.816362 | 118.0144577 | 2.235129101 | 132.406588 |
| 0.128332801 | 14.07041918 | 113.3475969 | 12513.59143 | 0.300791996 | 5003.866362 | 152.6044577 | 3.375129101 | 123.136588 |
| 0.098052939 | 13.63129945 | 119.0445425 | 24994.88836 | 0.357748332 | 3306.651559 | 92.33267529 | 3.010141536 | 127.7517707 |
| 0.108052939 | 16.94129945 | 120.3445425 | 21934.13836 | 0.357748332 | 3113.391559 | 94.62267529 | 3.170141536 | 202.0917707 |
| 0.188052939 | 14.42129945 | 350.7145425 | 21556.46836 | 0.827748332 | 3164.181559 | 72.70267529 | 2.280141536 | 207.4017707 |
| 0.078052939 | 20.99129945 | 124.4545425 | 24891.41836 | 0.477748332 | 3689.181559 | 92.91267529 | 3.030141536 | 687.8717707 |
| 0.090054474 | 15.32412942 | 123.9782162 | 22045.65109 | 0.384130732 | 4528.445075 | 99.9155753 | 3.185625507 | 151.603075 |
| 0.160054474 | 17.42412942 | 269.8982162 | 20863.61109 | 0.664130732 | 4052.165075 | 127.3255753 | 2.865625507 | 123.693075 |
| 0.100054474 | 16.28412942 | 106.8782162 | 19944.34109 | 0.354130732 | 4369.405075 | 132.6155753 | 3.585625507 | 172.183075 |
| 0.060054474 | 14.75412942 | 128.3182162 | 22957.22109 | 0.264130732 | 3989.225075 | 65.6255753 | 2.695625507 | 133.883075 |
| 0.098333903 | 14.28610826 | 114.1553261 | 19872.26769 | 0.37909072 | 4602.669928 | 131.6920903 | 2.916639972 | 235.9903987 |
| 0.118333903 | 14.42610826 | 122.5053261 | 24456.96769 | 0.24909072 | 3929.119928 | 125.2220903 | 3.226639972 | 164.1103987 |
| 0.078333903 | 11.58610826 | 173.2153261 | 19664.63769 | 0.39909072 | 4064.369928 | 84.7820903 | 1.956639972 | 179.3203987 |
| 0.088333903 | 14.81610826 | 126.8353261 | 21550.86769 | 0.75909072 | 5464.059928 | 174.0420903 | 3.186639972 | 488.5303987 |
| 0.098332801 | 13.34041918 | 157.9975969 | 20025.76143 | 0.350791996 | 4409.816362 | 105.7144577 | 3.235129101 | 139.266588 |
| 0.118332801 | 15.78041918 | 155.2375969 | 20584.19143 | 0.390791996 | 4063.806362 | 133.9744577 | 3.195129101 | 161.766588 |
| 0.078332801 | 13.85041918 | 125.2975969 | 24262.50143 | 0.300791996 | 5067.916362 | 104.9844577 | 3.345129101 | 145.816588 |

| Ni.CON | P.CON | Rb.CON | S.CON | Se.CON | Sr.CON | Zn.CON | Al.TOT | As.TOT |
|---|---|---|---|---|---|---|---|---|
| 0.473778303 | 1246.54791 | 6.417398035 | 1396.316312 | 0.467766417 | 64.93066088 | 29.10696278 | 0.320725794 | 0.002856076 |
| 0.343778303 | 1410.48791 | 7.627398035 | 1680.956312 | 0.397766417 | 58.01066088 | 17.79696278 | 0.178812092 | 0.002055478 |
| 0.323778303 | 1175.09791 | 6.007398035 | 1711.956312 | 0.467766417 | 73.95066088 | 33.52696278 | 0.250796985 | 0.0032411 |
| 0.263999868 | 1037.204032 | 4.510191986 | 1554.870086 | 0.552716313 | 68.14666522 | 14.77376592 | 0.16262074 | 0.002853178 |
| 0.263999868 | 1309.104032 | 10.24019199 | 1497.300086 | 0.492716313 | 47.72666522 | 7.943765923 | 0.254107428 | 0.003525575 |
| 0.223999868 | 1107.274032 | 4.980191986 | 1531.440086 | 0.602716313 | 57.78666522 | 30.07376592 | 0.17158974 | 0.003503751 |
| 0.389365345 | 1229.941274 | 9.090223394 | 1839.921039 | 0.427715473 | 59.84481832 | 20.12740932 | 0.248479871 | 0.002959057 |
| 0.289365345 | 1105.471274 | 6.940223394 | 1400.291039 | 0.457715473 | 56.94481832 | 14.65740932 | 0.253604819 | 0.003690015 |
| 0.289365345 | 1332.371274 | 10.70022339 | 1584.821039 | 0.477715473 | 56.02481832 | 16.26740932 | 0.525438348 | 0.004753082 |
| 0.24646484 | 1323.587524 | 7.088587798 | 1833.747537 | 0.505811799 | 74.94756708 | 26.27824854 | 0.202288201 | 0.004279108 |
| 0.18646484 | 1242.127524 | 9.748587798 | 1603.777537 | 0.425811799 | 53.94756708 | 24.26824854 | 0.280344494 | 0.005192583 |
| 0.40646484 | 907.8275237 | 6.458587798 | 1436.607537 | 0.475811799 | 71.05756708 | 21.65824854 | 0.620174088 | 0.004556622 |
| 0.103778303 | 654.3779098 | 11.80739804 | 1592.696312 | 0.647766417 | 32.01066088 | 17.95696278 | 0.160005634 | 0.001303216 |
| 0.213778303 | 646.0779098 | 10.45739804 | 1321.946312 | 0.717766417 | 36.01066088 | 53.06696278 | 0.136550977 | 0.00150623 |
| 0.453778303 | 594.7179098 | 11.93739804 | 1325.476312 | 0.717766417 | 28.79066088 | 16.52696278 | 0.482062312 | 0.000938944 |
| 0.733778303 | 652.2879098 | 12.12739804 | 1434.646312 | 0.747766417 | 38.35066088 | 25.50696278 | 0.139440319 | 0.001295475 |
| 0.323999868 | 507.3140318 | 11.79019199 | 1516.420086 | 0.622716313 | 45.03666522 | 14.83376592 | 0.129170033 | 0.001285115 |
| 0.543999868 | 606.1640318 | 9.620191986 | 1279.890086 | 0.752716313 | 45.76666522 | 26.40376592 | 0.340241533 | 0.001375204 |
| 0.373999868 | 576.1740318 | 8.580191986 | 1257.300086 | 0.692716313 | 49.46666522 | 61.38376592 | 0.11003696 | 0.001605644 |
| 0.303999868 | 564.5240318 | 9.880191986 | 1311.550086 | 0.572716313 | 39.94666522 | 12.25376592 | 0.145550735 | 0.001301901 |
| 0.329365345 | 458.5312738 | 9.930223394 | 1293.131039 | 0.627715473 | 51.81481832 | 25.72740932 | 0.103222213 | 0.001579372 |
| 0.419365345 | 585.8812738 | 9.910223394 | 1776.991039 | 0.727715473 | 40.57481832 | 21.64740932 | 0.138662696 | 0.001510602 |
| 0.349365345 | 526.9512738 | 9.520223394 | 1258.001039 | 0.737715473 | 43.08481832 | 23.25740932 | 0.162527407 | 0.000933733 |
| 0.339365345 | 496.2112738 | 8.820223394 | 1220.171039 | 0.697715473 | 69.38481832 | 25.45740932 | 0.122925161 | 0.001552937 |
| 0.22646484 | 497.2075237 | 8.358587798 | 1138.157537 | 0.675811799 | 49.32756708 | 26.01824854 | 0.023281846 | 0.000147397 |
| 0.26646484 | 546.8675237 | 8.368587798 | 1366.157537 | 0.665811799 | 44.76756708 | 25.90824854 | 0.013681407 | 0.000126041 |
| 0.49646484 | 607.1175237 | 10.6485878 | 2103.587537 | 0.775811799 | 52.11756708 | 25.88824854 | 0.030873428 | 0.000314399 |

| B.TOT | Ca.TOT | Cd.TOT | Co.TOT | Cu.TOT | Fe.TOT | K.TOT | Li.TOT | Mg.TOT |
|---|---|---|---|---|---|---|---|---|
| 0.538000776 | 45.28320508 | 0.001171319 | 0.000562961 | 0.04356122 | 0.425827665 | 43.14336616 | 0.001011094 | 16.73461125 |
| 0.60422909 | 32.39966289 | 0.0000938 | 0.000222936 | 0.039927914 | 0.278507383 | 29.46434357 | 0.000654234 | 11.77309198 |
| 0.729753771 | 58.22680236 | 0.000168866 | 0.00032295 | 0.043842508 | 0.426638979 | 41.79439283 | 0.000980673 | 18.89523655 |
| 0.528341085 | 39.45521881 | 0.0000676 | 0.000198042 | 0.041150163 | 0.296188365 | 25.10682444 | 0.001543999 | 13.55623129 |
| 0.806009041 | 42.44043984 | -0.0000796 | 0.00024616 | 0.04330501 | 0.384934826 | 56.70349602 | 0.000963251 | 14.19775574 |
| 0.759103876 | 49.10542968 | -0.00000911 | 0.000239855 | 0.053015171 | 0.356415562 | 35.29776671 | 0.000904341 | 17.13749711 |
| 0.886350907 | 51.27719792 | 0.000499506 | 0.000420636 | 0.050426743 | 0.443249063 | 60.58310606 | 0.001169804 | 18.64108102 |
| 1.148610742 | 51.8987404 | 0.00043926 | 0.000467733 | 0.046600974 | 0.39156487 | 43.21745475 | 0.000434045 | 18.06561486 |
| 0.999770483 | 65.07657443 | 0.000565807 | 0.000461644 | 0.060824479 | 0.63098685 | 78.82839054 | 0.00192054 | 24.95191167 |
| 0.737778918 | 61.74157589 | 0.000424325 | 0.000402564 | 0.059910017 | 0.421052944 | 47.31119021 | 0.001703676 | 21.52026938 |
| 1.042364252 | 66.94853928 | 0.000630128 | 0.000395735 | 0.076337298 | 0.554845097 | 94.69968398 | 0.001216473 | 21.5810637 |
| 1.037224292 | 48.25488256 | 0.000346708 | 0.000387817 | 0.042520304 | 0.34253239 | 37.81562638 | 0.000908983 | 15.12150544 |
| 0.428054971 | 13.66380252 | 0.0000863 | 0.000165085 | 0.022950078 | 0.200427077 | 42.0821677 | 0.000602316 | 5.567180914 |
| 0.385820955 | 18.95008324 | 0.0000728 | 0.000190566 | 0.0298782 | 0.21224336 | 38.68372533 | 0.000630936 | 5.490873721 |
| 0.27216091 | 9.249431514 | 0.0000817 | 0.000250793 | 0.019232691 | 0.467723769 | 28.74837343 | 0.001103911 | 4.219850467 |
| 0.324609309 | 15.04903099 | 0.0000858 | 0.000130632 | 0.035131688 | 0.208290975 | 41.65904822 | 0.000799574 | 6.174328447 |
| 0.540416353 | 16.33425004 | -0.000065 | 0.000137229 | 0.02335151 | 0.188922876 | 33.59402921 | 0.000585354 | 6.900622509 |
| 0.247577265 | 14.79083571 | 0.000614589 | 0.000215087 | 0.02341523 | 0.362699826 | 28.03734023 | 0.000892485 | 5.44545862 |
| 0.406268848 | 21.45515217 | 0.000120171 | 0.000155469 | 0.02530292 | 0.166071571 | 30.99030072 | 0.000550262 | 6.789353261 |
| 0.308152241 | 14.24542638 | 0.00024663 | 0.0000885 | 0.021745216 | 0.189120428 | 33.83525434 | 0.000389286 | 5.879476635 |
| 0.395802476 | 17.02268903 | 0.000182033 | 0.000137142 | 0.019924215 | 0.159207478 | 27.71498916 | 0.000528701 | 6.419143962 |
| 0.552525254 | 17.87242417 | 0.000200559 | 0.000214735 | 0.026178432 | 0.222305098 | 44.38099767 | 0.000452014 | 7.130003383 |
| 0.11894996 | 12.05095046 | 0.000135351 | 0.0000959 | 0.014189004 | 0.212129291 | 24.0824282 | 0.000488749 | 4.97745743 |
| 0.200592589 | 21.82276103 | 0.00015856 | 0.000126729 | 0.021256027 | 0.181965137 | 30.91809443 | 0.001089035 | 7.839049602 |
| 0.019086722 | 2.263393668 | 0.0000259 | 0.0000189 | 0.002560884 | 0.030329896 | 3.844230988 | 0.0000673 | 0.846527248 |
| 0.033995976 | 1.530074306 | 0.0000163 | 0.0000168 | 0.002240256 | 0.022038195 | 2.922220033 | 0.0000555 | 0.576915368 |
| 0.115109198 | 5.230716832 | 0.0000452 | 0.0000338 | 0.005982886 | 0.054124087 | 10.4805341 | 0.000129931 | 2.189158872 |

| Mn.TOT | Mo.TOT | Na.TOT | Ni.TOT | P.TOT | Rb.TOT | S.TOT | Se.TOT | Sr.TOT |
|---|---|---|---|---|---|---|---|---|
| 0.42043354 | 0.009190871 | 0.360079232 | 0.001418317 | 3.731704406 | 0.019211321 | 4.180055731 | 0.00140032 | 0.194378436 |
| 0.216683112 | 0.006230071 | 0.247375074 | 0.000781625 | 3.20692897 | 0.017341888 | 3.821874301 | 0.000904374 | 0.131894834 |
| 0.428016154 | 0.010737703 | 0.438684539 | 0.001066406 | 3.870338848 | 0.019786152 | 5.638552298 | 0.00154065 | 0.243566186 |
| 0.343333208 | 0.00966189 | 0.721230712 | 0.000653093 | 2.565876081 | 0.01115749 | 3.846498703 | 0.001367331 | 0.168583898 |
| 0.391215318 | 0.012212789 | 0.523994519 | 0.000927653 | 4.599981176 | 0.035982389 | 5.261271864 | 0.001731326 | 0.167703831 |
| 0.526091995 | 0.011146739 | 0.762773444 | 0.00076694 | 3.79112833 | 0.017051377 | 5.24340473 | 0.002063604 | 0.197852255 |
| 0.52584778 | 0.009870002 | 0.591318542 | 0.00138406 | 4.372018858 | 0.032312623 | 6.540287451 | 0.001520382 | 0.212727778 |
| 0.479462193 | 0.012598266 | 0.422490045 | 0.001054637 | 4.029063167 | 0.025294731 | 5.103579966 | 0.001668216 | 0.207544308 |
| 0.628624331 | 0.015570486 | 0.401911423 | 0.001358471 | 6.255025585 | 0.050233874 | 7.440190538 | 0.00224271 | 0.263017283 |
| 0.629990911 | 0.011754226 | 0.86895213 | 0.000838465 | 4.502797485 | 0.024115123 | 6.23834363 | 0.001720754 | 0.254968947 |
| 0.596204825 | 0.011291792 | 0.668913354 | 0.000942014 | 6.275183888 | 0.049249517 | 8.10222684 | 0.002151186 | 0.272541182 |
| 0.461165221 | 0.01019952 | 0.372114371 | 0.001228322 | 2.743422354 | 0.019517622 | 4.34137667 | 0.001437886 | 0.21473343 |
| 0.15545415 | 0.005067967 | 0.215086835 | 0.000174724 | 1.101730903 | 0.019879301 | 2.681512809 | 0.0010906 | 0.053894139 |
| 0.166879479 | 0.00559096 | 0.356415302 | 0.000377026 | 1.139442999 | 0.018442991 | 2.331425433 | 0.001265875 | 0.063509516 |
| 0.096958538 | 0.003040867 | 0.276597421 | 0.000605173 | 0.793134212 | 0.015920084 | 1.767696236 | 0.000957236 | 0.038396116 |
| 0.155501529 | 0.005071339 | 1.151243487 | 0.001228074 | 1.091689236 | 0.020296789 | 2.401068474 | 0.001251485 | 0.064184853 |
| 0.152255279 | 0.004854381 | 0.231018721 | 0.000493724 | 0.773065052 | 0.017966358 | 2.3107805 | 0.00094892 | 0.06862864 |
| 0.17110511 | 0.00385094 | 0.166223614 | 0.000731048 | 0.814587039 | 0.012927992 | 1.719966579 | 0.00101153 | 0.061503043 |
| 0.206063291 | 0.005571486 | 0.267544826 | 0.000581136 | 0.895281846 | 0.013332239 | 1.953642267 | 0.00107637 | 0.076863248 |
| 0.096721551 | 0.003972919 | 0.197322136 | 0.000448047 | 0.832017696 | 0.014561815 | 1.933014042 | 0.000844092 | 0.058874965 |
| 0.183665242 | 0.004067711 | 0.329125566 | 0.000459352 | 0.639493664 | 0.013849252 | 1.803473729 | 0.000875448 | 0.072263878 |
| 0.227235092 | 0.005855243 | 0.297804017 | 0.000761004 | 1.063173316 | 0.017983652 | 3.224628504 | 0.001320554 | 0.073629362 |
| 0.103828946 | 0.002396212 | 0.219605909 | 0.000427853 | 0.645334352 | 0.011659004 | 1.540619266 | 0.000903448 | 0.052764107 |
| 0.249690632 | 0.004571734 | 0.700873358 | 0.000486873 | 0.711892776 | 0.012653992 | 1.750526425 | 0.001000982 | 0.099543387 |
| 0.0202934 | 0.000621029 | 0.026734211 | 0.0000435 | 0.095446087 | 0.00160455 | 0.218485599 | 0.000129732 | 0.009469131 |
| 0.019019588 | 0.000453594 | 0.022965078 | 0.0000378 | 0.077635657 | 0.001188041 | 0.193945579 | 0.0000945 | 0.006355396 |
| 0.045349536 | 0.001444976 | 0.062987558 | 0.000214455 | 0.262253087 | 0.00459981 | 0.908674688 | 0.000335123 | 0.022512928 |

| Zn.TOT |
| ---: |
| 0.087135505 |
| 0.040463725 |
| 0.110425442 |
| 0.036547923 |
| 0.027913117 |
| 0.102967741 |
| 0.071546028 |
| 0.053421224 |
| 0.0763699 |
| 0.089397663 |
| 0.122602325 |
| 0.065450454 |
| 0.030232898 |
| 0.093590538 |
| 0.022040869 |
| 0.042689242 |
| 0.022604275 |
| 0.035482418 |
| 0.095380507 |
| 0.018060082 |
| 0.035880901 |
| 0.039282614 |
| 0.02848234 |
| 0.03652264 |
| 0.004994574 |
| 0.003678046 |
| 0.011182799 |
