## Supplemental Table 3 for "Inoculation with the mycorrhizal fungus *Rhizophagus irregularis* modulates the relationship between root growth and nutrient content in maize (*Zea mays* ssp. *mays* L.)"

| Trait | Units | B73.NC.mean | B73.M.mean | W22.NC.mean | W22.M.mean | adjP.fungus | adjP.line | adjP.fungusxline |
|---|---|---|---|---|---|---|---|---|
| Hyphae | % | 0.148 | 90.6 | 0 | 91.2 | 0 | 0.9 | 0.913 |
| Vesicles | % | 0 | 64.3 | 0 | 59.8 | 0 | 0.444 | 0.636 |
| Arbuscules | % | 0 | 65.7 | 0 | 61.8 | 0 | 0.404 | 0.584 |
| QLA5 | cm | 2.04 | 2.19 | 2.23 | 2.28 | 0.112 | 0.02 | 0.658 |
| QLA10 | cm | 3.61 | 3.57 | 3.75 | 3.54 | 0.559 | 0.756 | 0.819 |
| QLA15 | cm | 5.12 | 4.76 | 5.22 | 5.42 | 0.711 | 0.068 | 0.314 |
| QLA20 | cm | 6.42 | 5.88 | 5.94 | 6.75 | 0.559 | 0.531 | 0.095 |
| QLA25 | cm | 7.6 | 7.46 | 8.08 | 8.25 | 0.935 | 0.062 | 0.819 |
| QLA30 | cm | 8.47 | 8.46 | 9.15 | 10.2 | 0.096 | 0 | 0.279 |
| QLA35 | cm | 9.23 | 9.66 | 10.6 | 11.4 | 0.128 | 0 | 0.819 |
| QLA40 | cm | 9.94 | 10.6 | 11.4 | 13.9 | 0 | 0 | 0.253 |
| QLA45 | cm | 10.2 | 13.3 | 13.1 | 16.3 | 0 | 0 | 0.99 |
| QLA50 | cm | 10.7 | 16 | 13.7 | 18.7 | 0 | 0 | 0.89 |
| QLA55 | cm | 11.9 | 17.9 | 15.5 | 21.4 | 0 | 0 | 0.99 |
| SFW | g | 10.1 | 28.6 | 18.1 | 42.9 | 0 | 0 | 0.299 |
| RL | cm | 72.4 | 90.8 | 93.2 | 99.1 | 0.007 | 0.002 | 0.372 |
| RV | mL | 13.3 | 37.3 | 24.3 | 59.5 | 0 | 0 | 0.253 |
| RWF | g | 13 | 34.8 | 22.5 | 54.2 | 0 | 0 | 0.257 |
| SDW | g | 1.27 | 3.45 | 2.32 | 5.13 | 0 | 0 | 0.526 |
| RDW | g | 0.971 | 2.71 | 1.84 | 3.86 | 0 | 0 | 0.665 |
| ARW | cm | 0.0604 | 0.0612 | 0.0632 | 0.0616 | 0.827 | 0.328 | 0.687 |
| NB | NA | 1.62 | 1.38 | 1.56 | 1.36 | 0.002 | 0.557 | 0.913 |
| NCC | n | 0.994 | 1.08 | 1.36 | 1.02 | 0.174 | 0.042 | 0.133 |
| ND | cm | 44.7 | 52.2 | 53 | 57.4 | 0.009 | 0.004 | 0.716 |
| EAR | cm/cm | 0.308 | 0.349 | 0.322 | 0.298 | 0.69 | 0.425 | 0.279 |
| NLD | NA | 0.925 | 0.651 | 0.95 | 0.672 | 0 | 0.786 | 0.99 |
| MEA | cm | 42.7 | 49.1 | 48.6 | 58 | 0.002 | 0.004 | 0.751 |
| MaxNR | n | 31.6 | 50.9 | 41.2 | 57 | 0 | 0.004 | 0.74 |
| NW | cm | 15.3 | 20.9 | 21.3 | 22.4 | 0.007 | 0.002 | 0.261 |
| MNR | n | 20.1 | 37 | 27 | 42.6 | 0 | 0.002 | 0.913 |
| MinEA | cm | 13 | 16.5 | 15.4 | 17.1 | 0.003 | 0.066 | 0.545 |

| | | | | | | | | |
|---|---|---|---|---|---|---|---|---|
| NetA | cm2 | 75 | 159 | 117 | 202 | 0 | 0 | 0.99 |
| NCA | cm2 | 628 | 884 | 894 | 1070 | 0.002 | 0.002 | 0.797 |
| NP | cm | 2920 | 5970 | 4410 | 7780 | 0 | 0 | 0.896 |
| NS | NA | 0.126 | 0.182 | 0.133 | 0.188 | 0 | 0.419 | 0.99 |
| SRL | cm/cm3 | 298 | 285 | 283 | 287 | 0.69 | 0.538 | 0.665 |
| NSA | cm2 | 285 | 620 | 443 | 776 | 0 | 0 | 0.99 |
| NL | cm | 1500 | 3170 | 2270 | 4050 | 0 | 0.002 | 0.913 |
| NV | cm3 | 5.24 | 12.1 | 8.55 | 14.5 | 0 | 0.002 | 0.819 |
| NWD | NA | 0.341 | 0.406 | 0.403 | 0.395 | 0.205 | 0.219 | 0.279 |
| Crown | g | 2.01 | 4.21 | 2.6 | 5.24 | 0 | 0.002 | 0.636 |
| X15cm | g | 3.8 | 7.91 | 5.36 | 12.4 | 0 | 0 | 0.204 |
| X30cm | g | 3.35 | 5.64 | 5.28 | 9.77 | 0 | 0 | 0.314 |
| X45cm | g | 2.3 | 6.37 | 4.39 | 8.49 | 0 | 0 | 0.99 |
| X60cm | g | 0.915 | 5.21 | 2.46 | 6.75 | 0 | 0.005 | 0.994 |
| X75cm | g | 0.467 | 3.14 | 1.38 | 5.71 | 0 | 0.004 | 0.314 |
| X90cm | g | 0.142 | 2.33 | 0.984 | 5.83 | 0 | 0.017 | 0.279 |
| Root.PC1 | NA | 3.76 | -1.52 | 0.84 | -4.12 | 0 | 0 | 0.913 |
| Root.PC2 | NA | 0.292 | -0.332 | -0.475 | 0.742 | 0.601 | 0.892 | 0.261 |
| Root.PC3 | NA | 0.451 | -0.68 | 0.706 | -0.562 | 0.014 | 0.712 | 0.972 |
| Root.PC4 | NA | 0.277 | 0.388 | -0.756 | 0.0167 | 0.295 | 0.064 | 0.64 |
| Root.PC5 | NA | 0.308 | -0.0717 | -0.0443 | -0.278 | 0.334 | 0.382 | 0.913 |
| Root.PC6 | NA | 0.134 | 0.442 | -0.184 | -0.347 | 0.69 | 0 | 0.299 |
| Al.CON | ppm | 119 | 85.1 | 85.7 | 87.6 | 0.336 | 0.329 | 0.526 |
| As.CON | ppm | 0.872 | 1.06 | 1.16 | 1.69 | 0.002 | 0 | 0.279 |
| B.CON | ppm | 221 | 240 | 168 | 195 | 0.2 | 0.004 | 0.913 |
| Ca.CON | ppm | 10800 | 14900 | 8050 | 10900 | 0 | 0 | 0.526 |
| Cd.CON | ppm | 0.112 | 0.103 | 0.178 | 0.159 | 0.711 | 0.064 | 0.972 |
| Co.CON | ppm | 0.104 | 0.106 | 0.111 | 0.103 | 0.827 | 0.828 | 0.842 |
| Cu.CON | ppm | 15.2 | 14.7 | 11.5 | 12.5 | 0.827 | 0 | 0.616 |
| Fe.CON | ppm | 155 | 120 | 123 | 130 | 0.289 | 0.343 | 0.279 |
| K.CON | ppm | 22000 | 14000 | 26200 | 22700 | 0 | 0 | 0.133 |

| | | | | | | | | |
|---|---|---|---|---|---|---|---|---|
| Li.CON | ppm | 0.434 | 0.332 | 0.213 | 0.315 | 0.926 | 0.002 | 0.174 |
| Mg.CON | ppm | 4120 | 5180 | 4010 | 4060 | 0.005 | 0.005 | 0.133 |
| Mn.CON | ppm | 109 | 137 | 121 | 160 | 0 | 0.052 | 0.687 |
| Mo.CON | ppm | 2.99 | 3.19 | 4.1 | 4.21 | 0.448 | 0 | 0.931 |
| Na.CON | ppm | 221 | 159 | 158 | 162 | 0.331 | 0.296 | 0.526 |
| Ni.CON | ppm | 0.365 | 0.308 | 0.414 | 0.365 | 0.289 | 0.289 | 0.99 |
| P.CON | ppm | 568 | 1200 | 659 | 1110 | 0 | 0.804 | 0.133 |
| Rb.CON | ppm | 10.1 | 7.48 | 11.7 | 10.8 | 0.002 | 0 | 0.279 |
| S.CON | ppm | 1410 | 1590 | 1270 | 1250 | 0.191 | 0 | 0.276 |
| Se.CON | ppm | 0.692 | 0.479 | 0.632 | 0.516 | 0 | 0.542 | 0.257 |
| Sr.CON | ppm | 44.4 | 61.9 | 33.1 | 40.7 | 0 | 0 | 0.257 |
| Zn.CON | ppm | 26.8 | 21.4 | 28.3 | 29.9 | 0.596 | 0.165 | 0.526 |
| Al.TOT | mg | 0.149 | 0.289 | 0.205 | 0.449 | 0 | 0.038 | 0.526 |
| As.TOT | mg | 0.00112 | 0.00362 | 0.00303 | 0.00901 | 0 | 0 | 0.253 |
| B.TOT | mg | 0.29 | 0.818 | 0.397 | 1.01 | 0 | 0.081 | 0.797 |
| Ca.TOT | mg | 13.4 | 51 | 19.5 | 55.3 | 0 | 0.148 | 0.913 |
| Cd.TOT | mg | 0.000134 | 0.00036 | 0.000414 | 0.000835 | 0.006 | 0.002 | 0.653 |
| Co.TOT | mg | 0.000132 | 0.000361 | 0.000254 | 0.000527 | 0 | 0 | 0.792 |
| Cu.TOT | mg | 0.0196 | 0.0501 | 0.0271 | 0.0649 | 0 | 0.034 | 0.687 |
| Fe.TOT | mg | 0.192 | 0.413 | 0.286 | 0.669 | 0 | 0 | 0.279 |
| K.TOT | mg | 28.1 | 49.5 | 60.4 | 118 | 0 | 0 | 0.226 |
| Li.TOT | mg | 0.000558 | 0.00112 | 0.000452 | 0.00161 | 0 | 0.275 | 0.253 |
| Mg.TOT | mg | 5.1 | 17.7 | 9.16 | 20.5 | 0 | 0.002 | 0.791 |
| Mn.TOT | mg | 0.137 | 0.471 | 0.291 | 0.819 | 0 | 0 | 0.256 |
| Mo.TOT | mg | 0.00376 | 0.0109 | 0.00972 | 0.0222 | 0 | 0 | 0.279 |
| Na.TOT | mg | 0.301 | 0.532 | 0.327 | 0.86 | 0 | 0.087 | 0.279 |
| Ni.TOT | mg | 0.000471 | 0.00104 | 0.000945 | 0.00207 | 0.005 | 0.017 | 0.599 |
| P.TOT | mg | 0.729 | 4.16 | 1.46 | 5.65 | 0 | 0 | 0.336 |
| Rb.TOT | mg | 0.0131 | 0.0268 | 0.0267 | 0.0558 | 0 | 0 | 0.257 |
| S.TOT | mg | 1.78 | 5.48 | 2.95 | 6.33 | 0 | 0.007 | 0.842 |
| Se.TOT | mg | 0.000874 | 0.00165 | 0.00149 | 0.00268 | 0 | 0 | 0.548 |

| | | | | | | | | |
|---|---|---|---|---|---|---|---|---|
| Sr.TOT | mg | 0.0548 | 0.211 | 0.0801 | 0.206 | 0 | 0.413 | 0.526 |
| Zn.TOT | mg | 0.0347 | 0.0737 | 0.0655 | 0.152 | 0 | 0 | 0.261 |
